# vmrseq: probabilistic modeling of single-cell methylation heterogeneity

**DOI:** 10.1101/2023.11.20.567911

**Authors:** Ning Shen, Keegan Korthauer

## Abstract

Single-cell DNA methylation measurements reveal genome-scale inter-cellular epigenetic heterogeneity, but extreme sparsity and noise challenges rigorous analysis. Previous methods to detect variably methylated regions (VMRs) have relied on predefined regions or sliding windows, and report regions insensitive to heterogeneity level present in input. We present vmrseq, a statistical method that overcomes these challenges to detect VMRs with increased accuracy in synthetic benchmarks and improved feature selection in case studies. vmrseq also highlights context-dependent correlations between methylation and gene expression, supporting previous findings and facilitating novel hypotheses on epigenetic regulation. vmrseq is available at https://github.com/nshen7/vmrseq.

## Background

DNA methylation (DNAme) is an epigenetic modification that plays a crucial role in regulating gene expression and maintaining cellular identity in living organisms [1, 2]. Bisulfite sequencing (BS-seq) [3, 4] has become a widely-used technology to measure DNA methylation at a single-nucleotide resolution. Traditional bisulfite sequencing protocols, also referred to as ‘bulk’ BS-seq, allow for measurement of methylation level on a collection of cells. Though useful in many settings, bulk technology only quantifies the average signal seen in a population of cells that may consist of multiple cell types or states, each with unique methylation patterns. As a result, bulk BS-seq is not able to detect heterogeneity of inter-cellular methylation or effectively characterize cell identities. While cell type deconvolution algorithms [5–8] can estimate cell type compositions for bulk data, they either require reference databases of known cell types or offer a limited level of resolution and reliability.

To overcome the limitation of bulk technologies, protocols for methylation sequencing at single-cell resolution such as scBS-seq [9] (Fig. 1a) have been developed. These protocols have shown that DNAme can be an accurate marker distinguishing individual cells under different conditions or cell types. However, due to the small amounts of input genomic DNA in single cells and the destructive nature of bisulfite treatment, these technologies are limited by the sparsity and noisiness of the output data. Typically the vast majority of CpG dinucleotides are not observed (ranging from approximately 80% to 95+% in high-throughput studies) [10]. Additional sources of noise in single-cell data, as compared to bulk data, include increased technical variability due to amplification applied on limited amounts of materials, which tends to be uneven, biased, and error-prone [11].

**Fig. 1.**
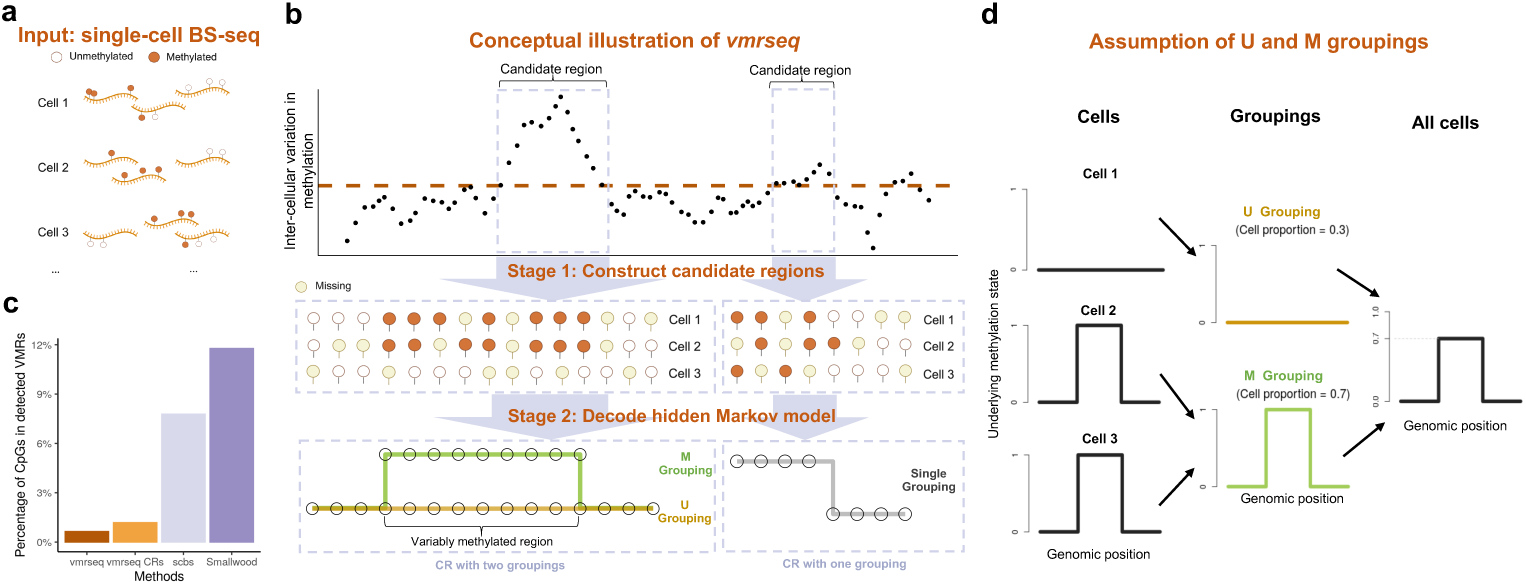
Overview of vmrseq framework. **a** vmrseq takes processed and filtered single-cell bisulfite sequencing binary methylation values as input. After processing and filtering, each sequenced CpG takes a value of methylated or unmethylated in each cell. Missing data indicates a lack of read coverage. **b** Detecting heterogeneity in single-cell methylation with vmrseq. In brief, vmrseq first defines candidate regions as those with consecutive CpG sites exhibiting cell-to-cell variation in methylation levels above a threshold that represents significantly high variance under a null condition; then vmrseq detects variably methylated regions by decoding one- and two-group hidden Markov models fit on sites within candidate regions. **c** VMRs from vmrseq generated the fewest false positives in comparison to CRs and other methods. y-axis shows the percentage of variably methylated CpGs out of all covered sites from a homogeneous cell population. Cells from the subtype *mL4* (n=370) in the annotated single-cell bisulfite sequencing dataset published by Luo et al. [27] were selected to establish a homogeneous cell population, which served as the input for the analytical methods. **d** vmrseq relies on the assumption of M/U groupings. In short, cells can be divided into an M grouping and a U grouping according to their underlying methylation states, assuming each CR holds at most one VMR and every cell exhibits uniform hidden states within the VMR if any, i.e., they are either all methylated or all unmethylated.

Assessing the cellular heterogeneity of DNAme is already challenging in the presence of noise and biases intrinsic to single-cell technologies. In addition, a considerable number of studies have stressed the existence of spatial correlations of DNA methylation across nearby loci; this correlation implies that individual CpGs are not likely to impact epigenetic function on their own but rather through biochemical interactions with several loci together [12–14]. Variable methylation exhibited by individual loci might be more likely to arise from technical noise. Moreover, many loci-level discoveries may originate from a single regional discovery hence should not be counted multiple times.

To reflect these groupings of CpGs, identifying regions with distinctive methylation levels across cells, referred to as variably methylated regions (VMRs) [15, 16], is considered one of the main analytical objectives in the analysis of scBS-seq data. The identified variable regions may serve as epigenetic features of cell types and states and facilitate integrative analyses of single-cell multi-omics assays. They might also foster understanding of environmental influence [17, 18], allowing the identification of epigenetic changes in response to extrinsic stimuli. However, defining the CpG clusters is a challenging task in itself since they may occur anywhere on the genome with diverse sizes and in various location contexts.

Efforts have been made for efficient VMR selection and clustering inference through non-probabilistic methods [19–22]. Smallwood et al. [9] and scbs [23] both rank sliding windows by cell-to-cell variances of mean methylation levels, but use different statistics to measure variance. A number of probabilistic methods have also been proposed. Melissa [24] and Epiclomal [25] propose to directly infer cell clusters through probabilistic graphical models hence are not presented as feature selection tools. scMET [26] models cell-to-cell heterogeneity through a hierarchical Bayesian model and selects features through ranking a statistic that represents heterogeneity.

All of these feature selection methods share a common drawback that features are selected from a set of pre-defined genomic regions, for example, gene promoters or sliding windows. Such analytical approaches only provide region-level resolution in the analysis since they operate on aggregated counts of the pre-defined regions without considering the case where VMRs exist outside the regions or overlap with edge of region boundaries. While the settings for window width and step size in scbs offer some flexibility, they have not reached the level of achieving base pair-level resolution. We reasonably hypothesize that these regions should not be restricted to genomic ranges specified a priori.

To address the limitations and accurately pinpoint inter-cellular heterogeneity from single-cell DNAme data, here we present *vmrseq*, a statistical approach to accurately and robustly detect VMRs without the need for prior knowledge of their sizes or location contexts (Fig. 1b). Our results both on simulated and previously published experimental datasets demonstrate that it outperforms existing methods for detecting and quantifying DNAme heterogeneity. The reanalysis of two recent studies involving single-cell bisulfite sequencing data reveals that vmrseq identifies biologically relevant regions with high variability across cells, leading to significantly enhanced cell clustering performance. Moreover, vmrseq highlights context-dependent correlation patterns between gene expression and DNAme that support previous findings and may inform new biological hypotheses regarding the involvement of epigenetic variability in the cell cycle.

## Results

### Pinpoint cell-to-cell DNAm heterogeneity with vmrseq

vmrseq is a two-stage approach that first constructs candidate regions (CRs) and then determines whether a VMR is present and its location if applicable. The input to vmrseq is a matrix of binary methylation values where each row is a CpG site and each column is an individual cell (Fig. 1a). To avoid ambiguities, sites with intermediate methylation level between 0 and 1 are filtered out for each cell.

Stage 1 of vmrseq (Fig. 1b) scans the genome for regions containing consecutive CpGs that show evidence of potential cell-to-cell variation (i.e., CRs). As the methylation levels of neighboring CpGs display strong correlation, vmrseq first uses smoothing to mitigate the influence of limited coverage and counteract the reduction in statistical power caused by the inherent noise in single-cell data.

Specifically, the candidate regions are constructed by first applying a kernel smoother to ‘relative’ methylation levels of individual cells that are in reference to across-cell average methylation on CpG sites (“Methods”). Next, groups of consecutive loci that exceed some threshold on the variance of smoothed relative methylation levels are selected. Such a threshold can be computed by taking the 1 − *α* quantile of an approximate null distribution of variance, where *α* is the designated significance level. This distribution of variance is simulated from labeled data while taking the size of input dataset into account (“Methods” and Additional File 1: Section S1.2). Such strategy enables control of false positives, showed by substantially reduced number of detected sites with variable methylation from a homogeneous dataset, compared to other methods (Fig. 1c).

The stage 2 of vmrseq (Fig. 1b) optimizes a hidden Markov model (HMM) that models methylation states of individual CpG sites for each CR (“Methods”). To be more specific, for each cell, we assume every CpG site has an unobserved methylation state, modeled as a binary hidden state, where 1 represents methylated and 0 unmethylated. The observed methylation level from bisulfite sequencing is assumed to be determined by both the hidden state and technical error. The estimation of parameters and hidden states in the HMM determines whether groups of cell subpopulations show distinct epigenetic signals in each region and solves for the precise genomic range of VMRs.

Since single-cell data usually contains a large and unknown number of cell subpopulations, we make a critical assumption of the existence of unmethylated (U) and methylated (M) groupings (Fig. 1d) to reduce model complexity and ease computational burden. Specifically, we assume that each CR contains at most one VMR, and every cell has uniform hidden states (i.e., all methylated or all unmethylated) in the VMR if any. Under this assumption, if cells are heterogeneous in terms of underlying states within a CR, then they can be partitioned into two groupings (referred to as the U grouping and M grouping) based on their estimated hidden states within the VMR. This partition remains applicable irrespective of the overall number of cell subpopulations, which is not known or inferred by the model. That is to say, we may infer the existence of VMRs by detecting the presence of the two groupings. On the other hand, if the cells are homogeneous, all cells should have identical sequences of hidden states across CpGs in this CR.

Therefore, to determine whether both U and M groupings exist for each CR, a one-state and a two-state HMM are optimized for single-grouping and two-grouping assumptions respectively, followed by decoding the corresponding hidden states. HMMs are adopted as they can model spatial correlation between CpG sites with transitions between different hidden states and effectively handle noisy data through modeling the emission probability with a hierarchical probabilistic structure. Subsequently, we may infer the presence of one or two groupings by comparing the maximum likelihood of the two models. This comparison of one- and two-grouping likelihood resembles the idea of statistical hypothesis testing, where one-grouping case is considered the null hypothesis and is rejected if two-grouping likelihood surpasses one-grouping. However, we have not developed formal p-value quantification due to the biases of CRs towards high variation and the lack of strictly nested HMMs (detailed in “Discussion”). Finally, in the event that the presence of two groupings are deemed more likely than single grouping, vmrseq delineates the boundary of a VMR by removing any CpGs with estimates of hidden states uniform across the two groupings, effectively acting as a trimming step due to the assumption of at most one VMR per CR. Evaluation with and without this trimming step are included in the following sections, and a detailed description of the methodology is provided in the “Methods” section. vmrseq is implemented as an R package and is freely available at https://github.com/nshen7/vmrseq.

### vmrseq improves accuracy in detecting heterogeneity in synthetic datasets

To benchmark the performance of vmrseq and alternative methods, synthetic data were constructed by adding simulated VMRs to scBS-seq data of chromosome 1 in a homogeneous cell population (with the assumption that they contain no VMRs). A wide array of simulation settings were included to evaluate the methods across diverse attributes of input data, including number of cells, sparsity level and number of cell subpopulations that show distinctive methylation profiles (see “Methods” for details about the simulation settings). The three levels of sparsity, determined by stratifying and subsampling cells in experimental data, represent on average 94.9% (high), 92.8% (medium) and 90.4%(low) unobserved CpGs in a cell. 2000 VMRs of size between 5 and 500 CpGs (600 bp *<* width *<* 26,000 bp; Additional File 1: Fig. S1), containing roughly 5% of total number of CpGs in the chromosome, were added to each dataset in order to evaluate precision and recall.

Four evaluation metrics were used: precision, recall, F1 score and ratio of relative areas (RRA) of the precision-recall curve, where RRA is similar to area under curve but restricted to a region of interest (0.8 ≤ precision ≤ 1 in our case; see “Methods” for more details). These metrics are computed for two manners of defining a true positive: 1) an individual CpG correctly detected in VMRs, referred as ‘CpG-based’; and 2) an individual detected region with least 3 sites overlap with any true VMR, referred as ‘region-based’.

For each simulation, we assessed the performance of vmrseq, stage 1 only of vmrseq (denoted as ‘vmrseq CRs’), and three other previously published methods that also aim to search genome-wide for regions showing inter-cellular variation in methylation as measured by scBS-seq data: scbs [23], Smallwood et al. [9] (denoted as ‘Smallwood’) and scMET [26]. See “Methods” for details about implementation and parameter settings of the methods.

In general, all methods except scbs seem to benefit from lower sparsity, and in most cases the number of cells in input did not have strong influence on method performance (Fig. 2 and Additional File 1: Fig. S2). It may appear that a larger size of sliding windows (i.e. 3kb used by Smallwood and 20kb used by scMET) results in reduced performance compared to a smaller smoothing bandwidth or window size (i.e. 2kb bandwidth used by vmrseq and 2kb windows used by scbs). However, we conducted an additional set of experiments on both synthetic datasets and the mouse frontal cortex dataset showing that a smaller window size alone does not seem to meaningfully improve the accuracy of detecting VMRs (see Additional File 1: Figures 24-28 and Section S2.1 for a detailed discussion).

**Fig. 2.**
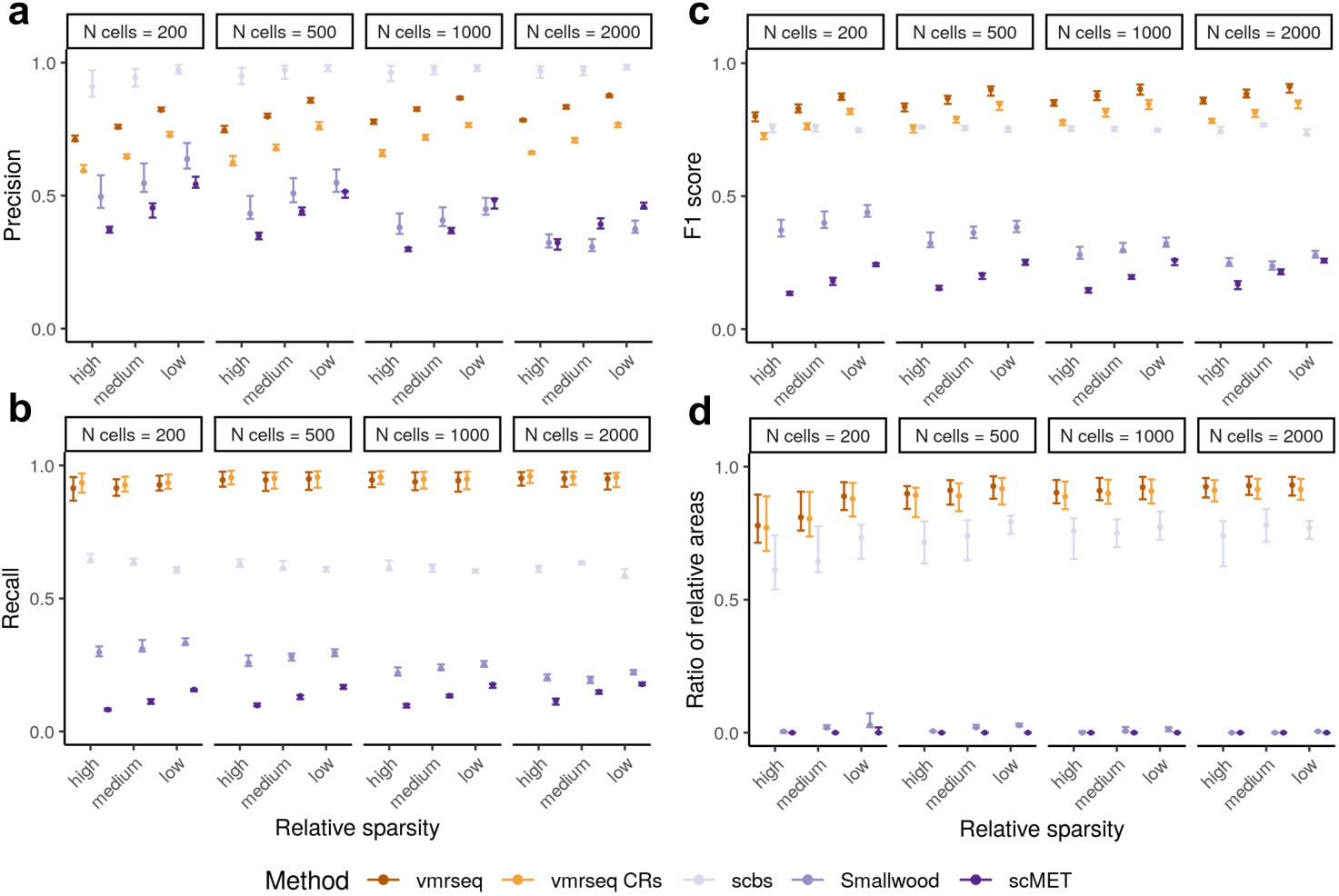
Region-based metrics evaluated on simulated VMRs, including **a** precision, **b** recall, **c** F1 score and **d** ratio of relative areas (“Methods”). A region-level true positive is defined as at least 3 sites overlap between detected and true VMR. Each interval consists of points originated from different number of subpopulations. Dot and boundaries of each interval indicates the maximum, median and minimum value of metric. Recall, precision and F1 score are computed using default parameter setting in each method.

In terms of comprehensive metrics such as F1 score and RRA, vmrseq rendered higher scores compared to other methods, suggesting that vmrseq is more accurate overall in detecting VMRs. In terms of specialized metrics such as precision and recall, candidate regions were able to achieve similar recall as vmrseq but less precision, indicating that stage 2 of vmrseq (i.e. decoding the HMM) effectively removed false positives from CRs. This effect of the HMM model was also shown in the homogeneous study in Fig. 1d. scbs achieved higher precision than vmrseq in these settings, but this came at the cost of a substantially lower recall. Presumably this is due to scbs using an arbitrary ad hoc threshold (top 2% variably methylated windows) to determine VMRs, which does not correctly reflect the level of heterogeneity present in the data. Note that we kept the 2% cutoff recommended by original paper because in general the amount of heterogeneity is not known in an experimental study.

We also conducted a second set of experiments with synthetic chromosomes wherein methylation levels are entirely simulated from the HMM model to ensure a firmly known null background. As expected, such simulation contained considerably less noise compared to the simulations based on experimental data. scMET failed to produce results in this simulation due to error evaluating probability at the initial value, hence only four methods were available for evaluation. We observed similar results from this set of simulations (Additional File 1: Fig. S3-S4). However, difference between CRs and vmrseq has reduced, suggesting that the full methodology of vmrseq is more suitable for noisy scBS-seq data, compared to stage 1 of vmrseq only.

### vmrseq enhances feature selection for single-cell methylomic unsupervised analysis

Although the primary objective of our proposed method is not to select features that optimize clustering performance, VMRs detected by vmrseq render reliable cell clusters. We applied vmrseq and the other methods to a dataset of 3,069 single-cell methylomes from mouse frontal cortex [27] to assess their efficacy in unsupervised cell clustering. This collection of cells spans 2 broad neuronal classes (i.e., excitatory and inhibitory) and 15 subtypes within those two classes (Fig. 3a). The original study annotated these groupings based on gene body non-CG methylation depletion in neuronal marker genes, making them suitable as a benchmark for clustering analyses. We applied vmrseq on this dataset along with the same competing methods that were evaluated in the simulation study, using their default parameters. In general, though vmrseq finds moderate number of VMRs compared to the other methods (Additional File 1: Fig. S5a), it tends to identify smaller regions and hence finds the fewest CpGs in detected regions (Additional File 1: Fig. S5b-d).

**Fig. 3.**
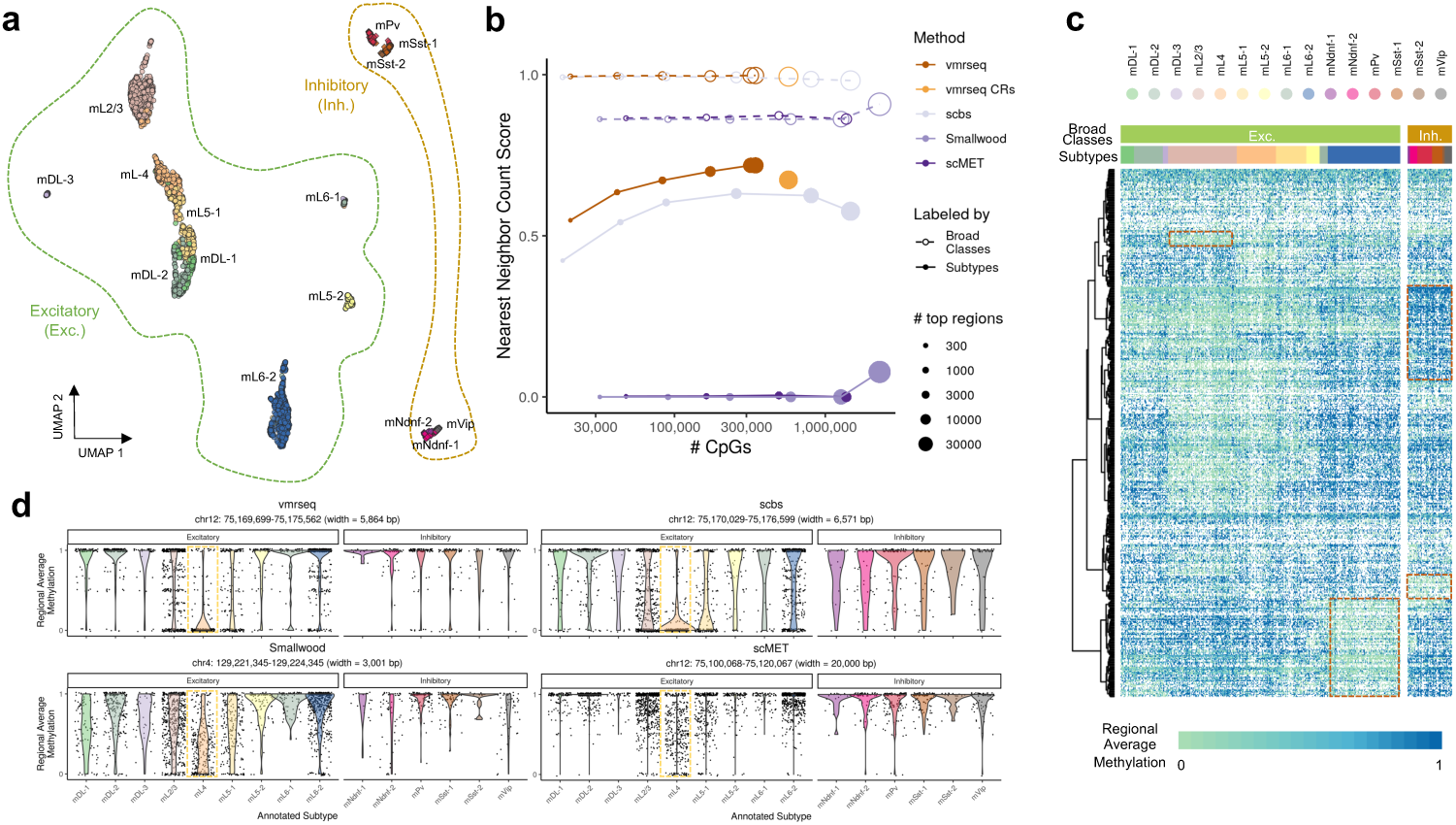
vmrseq outperforms other methods in unsupervised clustering analysis of mouse frontal cortex data. **a** vmrseq’s UMAP visualization of neuron subpopulations annotated in the mouse frontal cortex dataset. Coordinates were computed from regional average methylation levels of VMRs. Broad cell classes are indicated by dashed outlines; subtypes are indicated by colored points and labeled by text. **b** Evaluation of clustering performance in terms of nearest neighbor count score. The x-axis is the number of CpG sites in varying numbers of top-ranked VMRs (log-scaled). We use number of CpGs instead of number of regions as x-axis because the size of detected regions varies significantly across methods (Additional File 1: Fig. S5d). Regions were ranked by metrics proposed in each method respectively (“Methods”). The dot size indicates number of included regions. The top 300, 1000, 3000, 10000, and 30000 regions (if applicable) and all selected regions were extracted from each method respectively for computing the score. vmrseq CRs do not have rank thus only represented by one dot. Line and point types distinguish granularity of the cell type labels. The score, ranging from 0 to 1, quantitatively evaluates the quality of clustering by averaging the proportions of neighbors that share the same label (see “Methods” for details). **c** Heatmap of regional average methylation level of top-ranked 500 VMRs from vmrseq. Rows are sorted by hierarchical clustering; dashed red squares are examples of potential cell-type-specific marker regions; white color indicates missing. **d** Marker regions for cell type *mL4* detected by vmrseq exhibits more disparity of regional methylation level between the target and background cell types, compared to alternative methods. Annotated cell type labels from Luo et al. [27] were used to determine marker regions. Specifically, marker regions of cell type *mL4* among detected regions of each method are defined as those with absolute difference *>* 0.2 between the average methylation level of targeted cell type and all other cell types. Distribution of regional average methylation are plotted in violin shapes against cell subtypes; points represent methylation levels in individual cells.

To qualitatively evaluate the ability of each method to identify heterogeneity that distinguishes annotated cell types, we visualized the cells in a low-dimensional space by applying UMAP [28] on cell-to-cell dissimilarity matrices computed from regional average methylation of the detected regions (Fig. 3a for vmrseq, Additional File 1: Fig. S6 for others; “Methods”). Based on these UMAPs we may observe that vmrseq, vmrseq CRs and scbs seem to be markedly superior to the other two approaches in terms of unsupervised clustering analysis, where Smallwood and scMET were only able to identify the two broad classes as opposed to subtypes. We also note that Smallwood and scMET tended to favor genic regions and CpG islands, while vmrseq and scbs tended to select areas not only in genic regions and CpG islands but also intergenic and non-island regions (Additional File 1: Fig. S7-S10; “Methods”).

Further, to quantitatively assess the performance of feature selection, we employed a metric called ‘nearest neighbor count score’ [23]. This metric quantifies the quality of clustering by averaging the proportions of neighbors that share the same label (“Methods”). The scores were evaluated directly from the cell-to-cell dissimilarity matrix to avoid involving stochasticity introduced by dimension-reduction techniques or cluster partition methods, hence being a straightforward metric representing the degree to which the detected regions can distinguish cell types. We computed the scores in reference to the two sets of aforementioned cell type labels: the broad classes and the subtypes.

Fig. 3b plots the nearest neighbor count scores against the number of CpG sites in top-ranked regions. Notably, vmrseq identified significantly fewer CpG sites in detected regions compared to other methods, while performing better or equally well. In line with our observations from the UMAP figures (Fig. 3a, Additional File 1: Fig. S6), scMET and Smallwood effectively discriminated the two broad classes but were not able to separate subtypes within these classes. On the other hand, vmrseq, vmrseq CRs and scbs achieved nearly perfect nearest neighbor count scores (i.e., 1) in clustering the broad classes, and also performed commendably in distinguishing the subtypes. Similar conclusions were made based on evaluations using an alternative metric on clustering performance, the Silhouette scores [29] (see Additional File 1: Section S2.2 for a detailed discussion). In addition to genome-wide VMRs, we evaluated the clustering performance of VMRs that overlap with specific features such as histone modifications and gene promoters (Additional File 2: Table S1). All feature types achieved a perfect nearest neighbor count score for broad class labels. For the subtypes, distal H3K27ac and H3K4me1 peaks demonstrated superior clustering performance compared to gene promoters, however, none of the targeted regions achieved a score as high as that obtained by using all VMRs.

Moreover, we conducted experiments contrasting the region selection methods to a baseline approach where no region selection was performed and principal component analysis (PCA) is used on genome-wide large-sized bins to perform dimensionality reduction, similarly as proposed in SINBAD [22]. To ensure a fair comparison, PCA was also applied to the regions detected by each method. We observe that incorporating a dimension reduction step results in all methods achieving clustering performance comparable to that obtained using genome-wide information, despite the considerably smaller number of CpGs included in the detected regions (i.e., *<*350K CpGs for vmrseq and *>*1 million for all other methods). For a detailed discussion, please refer to Additional File 1: Section S2.3 [30, 31].

Additionally, vmrseq appears to exhibit a more favorable performance in the context of capturing cell type marker regions compared to the alternative methods. VMRs from vmrseq exhibited higher inter-cell-type variance compared to other methods both visually (Fig. 3c, Additional File 1: Fig. S11) and numerically (Additional File 1: Fig. S12). Specifically, some potential marker regions for broad classes or subtypes were identified visually from the display of 500 top-ranked regions (red dashed boxes in Fig. 3c). Top regions from scbs also seemed to contain some, but these were largely subject to sparsity (Additional File 1: Fig. S11a). Meanwhile, top regions from Smallwood and scMET did not show cell type-specific signals but instead seemed saturated with fully unmethylated and methylated regions respectively (Additional File 1: Fig. S11b-c). Fig. 3d shows the top-ranked hypomethylated cell type marker region for *mL4* (“Methods”) from the four methods respectively, where vmrseq displayed more extreme difference between the mean methylation of targeted cell type and of the others in background (vmrseq: diff = 0.62; scbs: 0.50; Smallwood: 0.37; scMET: 0.33).

### vmrseq captures heterogeneity associated with embryonic development and cell cycle states

In a second case study, we reanalyzed a single-cell multi-omics dataset of 939 cells profiled by scNMT-seq [32] (“Methods”). This dataset covers four early developmental stages of mouse embryos (E4.5-E7.5) and offers insights into dynamics of chromatin accessibility, DNA methylation, and RNA expression during the onset of gastrulation. We applied vmrseq on the single-cell DNAme data to pinpoint where methylation heterogeneity is exhibited across the genome. UMAP coordinates were obtained for each cell computed solely based on the methylation level of VMRs output by vmrseq (“Methods”). We did not apply other methods on this dataset due to its prior analysis in the original publications [23, 26], and its lack of ground truth for assessing methodological performance.

As found in the original study [32], there is a notable shift in the global methylation level within cells as developmental time progresses (Additional File 1: Fig. S13a). This led to heterogeneity observed across a substantial portion of the genome, resulting in the identification of 205,584 VMRs by vmrseq, which encompass approximately 28% of all CpGs in the mouse genome. This global shift also resulted in two distinctly separated groups of cells in the low-dimensional representation (Additional File 1: Fig. S13b-c).

It is important to note that the low global methylation level of early-stage lineages does not align with the model assumptions of vmrseq regarding the transition probability of switching hidden states. In particular, the transition probability trained from mature tissues is considered constant across cell subpopulations at a given CpG-CpG distance, and reaches to a plateau at large distances of around 80% global methylation level (or 1 - global methylation level ≈ 20%; Additional File 1: Fig. S14). However, despite such violation, we still made interesting discoveries from the regions detected by vmrseq.

Apart from two prominently separated clusters due to the shift in global methylation level over developmental time, we noticed that the second major source of heterogeneity uncovered by vmrseq was associated with different time points of the embryonic developmental stage (Fig. 4a). Stage E6.5 and E7.5 are closely related, aligning with previous discoveries of gene expression and chromatin accessibility from Argelaguet et al. [32].

**Fig. 4.**
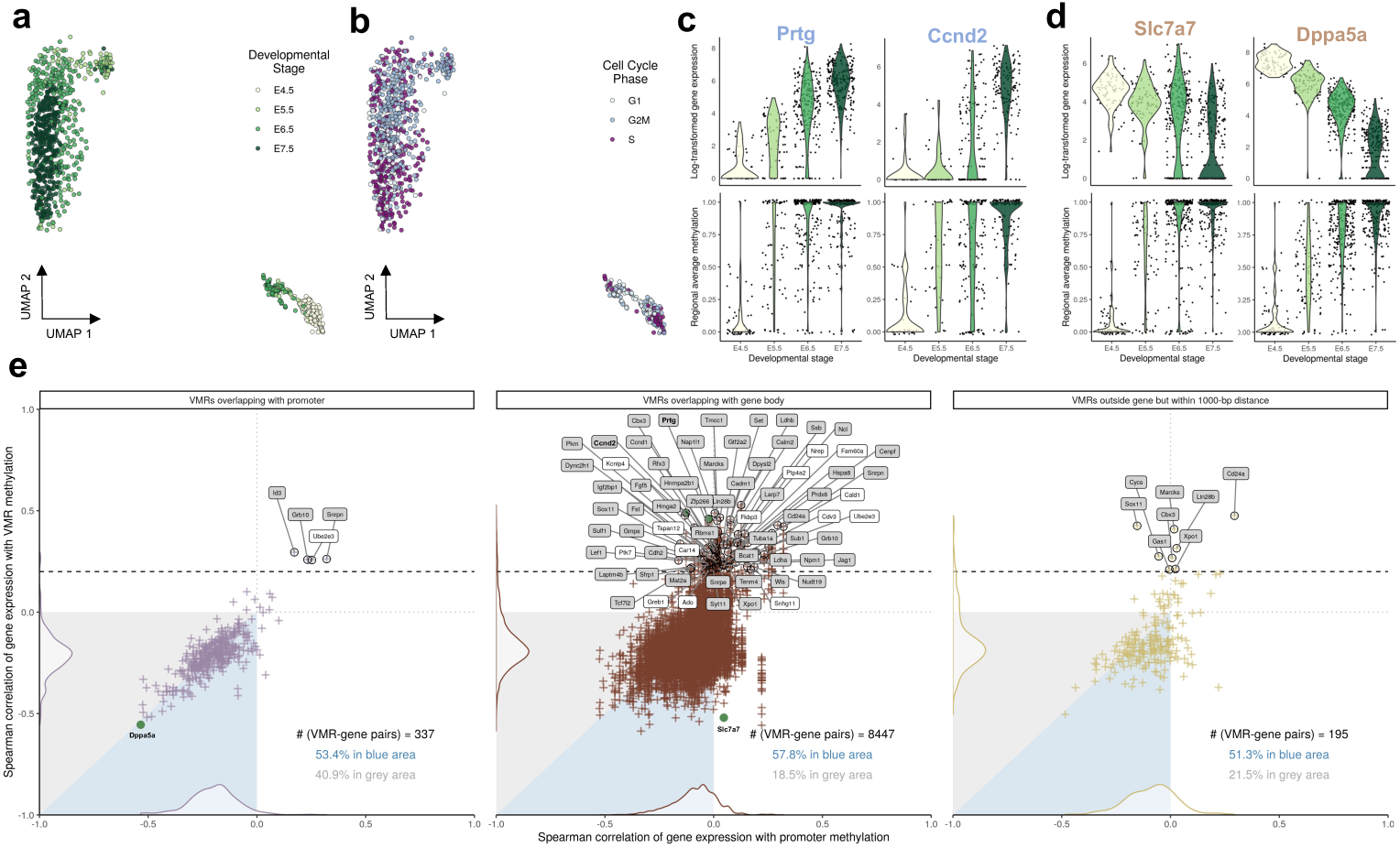
vmrseq applied to the multi-omics scNMT-seq gastrulation dataset captures heterogeneity associated with cell conditions and reveals complex associations between VMR methylation and RNA expression. **a, b** UMAP representation derived from regional average methylation levels of VMRs output by vmrseq, colored by developmental stage and cell cycle phase respectively. **c, d** Example genes whose expression levels are highly correlated with nearby VMR in positive and negative directions respectively. The x-axis displays developmental stage; the y-axis shows gene expression in log(x+1) scale (upper panels) and regional average methylation (lower panels). Each point represents an individual cell. **e** Comparison between Spearman correlation of gene expression with nearby VMRs methylation and that of gene expression with corresponding promoter methylation. Each point represents one VMR. Panels are partitioned by VMR’s position relative to the corresponding gene. Areas in blue represent where VMRs surpasses promoter in terms of absolute correlation with gene expression, and vice versa for grey areas. Black dashed line represents the threshold for highly positive correlation (cor *≥* 0.2). Symbols of genes exceeding this threshold are annotated in boxes. Gene set enrichment analysis were conducted for annotated genes and boxes with grey background color indicate genes in the enriched pathways (see Additional File 1: Fig. S16b-c for the pathways).

Given previous debate about whether DNAme varies throughout the cell cycle [33–35], we annotated the cell cycle phases based on solely gene expression (Fig. 4b, Additional File 1: S15b; “Methods”). Surprisingly, the VMRs seemed to contain some variation associated with cell cycle, where the cells in G2M phase displayed a more pronounced clustering tendency than the other two phases. Though cell cycle state is not observed as a primary driving factor, this suggests that the coupling between DNAme and cell cycle phases might be more intricate then previously acknowledged and deserve further investigation. In addition, we investigated the proportion of intermediate methylation values (i.e., between 0 and 1) in S-phase cells. It is reasonable to anticipate an increase in the proportion during the S phase of the cell cycle, reflecting DNA methylation maintenance. However, our analysis did not reveal a significantly higher proportion compared to the other two phases.

### vmrseq reveals bi-directional correlation between transcriptional and DNAme levels

Owing to the multi-modal nature of the scNMT-seq dataset [32], a valuable opportunity emerges for in-depth investigation into the linkage between inter-cellular variation in DNA methylation and transcription. Here, we explored correlation of gene expression to the methylation level of nearby VMRs as well as of corresponding promoters (Fig. 4c-e). In order to ensure meaningful correlations, we restricted our analysis to the genes with top 10% variable expression levels after normalizing with variance stabilization transformation [36] (“Methods”). For each highly variable gene, we linked VMRs within 1000-bp distance to that specific gene and assigned them into three mutually exclusive categories according to their relative position with respect to the gene: overlapping with promoter, overlapping with gene body and outside the gene (“Methods”).

Fig. 4e presents a comparison between the Spearman correlation of gene expression with linked VMRs and that of gene expression with corresponding promoters. Each cross-shaped dot in the figure represents a VMR-gene pair. We first observed that, in the left-bottom quadrant, there are substantially more dots in the blue triangles comparing with grey ones. This indicates that methylation of VMRs might be a stronger predictor of transcriptional activity in contrast with promoters, since they show higher correlation with the gene expression level.

Further, while promoters predominantly demonstrated negative or near-zero correlations, VMRs in gene bodies and outside genes demonstrated both positive and negative correlation with gene expression (Fig. 4e). Specifically, only 0.7% promoters exceeded 0.2 in terms of Spearman correlation with gene expression, whilst a more substantial proportion of VMRs in gene body (3%) and outside gene (4.1%) exhibited correlation greater than 0.2. For both directions, from the top ten ranked gene-VMR pairs with the highest absolute correlation, we selected two whose VMRs are ranked highest in terms of increment in log-likelihood of two-grouping model compared to one-grouping, and display them in Fig. 4c-d. Such distributions of correlation may imply the existence of two modes of regulation, supporting previous findings of positive correlation between gene body methylation and expression level [37–39]. Interestingly, we discovered that the genes highly positively correlated with VMRs (corr ≥ 0.2, labeled in boxes in Fig. 4e) are mostly significantly enriched in pathways relevant to RNA processing, gene regulation, and even regulation of cell cycle, suggesting an interplay between DNAme and various cellular processes (Additional File 1: Fig. S16b-c; “Methods”).

## Discussion

Single-cell DNAme sequencing assays can readily profile thousands of methylomes, spurring the need for methods to study cell-to-cell epigenomic heterogeneity across the entire genome, in addition to specific genic or CpG contexts. To address the challenges of technical noise and sparsity intrinsic to single-cell technologies, we introduced vmrseq, a statistical framework useful for detecting and prioritizing VMRs. In contrast to earlier methods that first define genomic windows for selection and then identify variation in mean methylation levels [9, 23, 26], vmrseq performs a genome-wide and context-free scan and directly models CpG sites hence accurately identifies VMRs at base pair-level resolution.

Through simulation studies, we showed that vmrseq is adaptable to a wide range of scenarios (Fig. 2). When applied to the mouse frontal cortex dataset, we observed that regions detected by vmrseq are more effective than those of alternative methods in distinguishing annotated cell types (Fig. 3a-b). As opposed to methods that use an arbitrary threshold for region selection [9, 23], our mechanism of candidate region construction and probabilistic modeling fosters region selection in accordance with the level of heterogeneity present in the input datasets. Furthermore, by revisiting the multi-omic mouse gastrulation dataset, our method detected variation sourced from embryonic development and cell cycle phases (Fig. 4a-b). Leveraging the multi-modal nature of the dataset, we found a set of genes that exhibited a positive correlation with the methylation of VMRs but not promoters, elucidating complex patterns of epigenetic regulation (Fig. 4c-e). Noteworthily, though only bisulfite sequenced samples have been included in the case studies, we believe that, with properly trained parameters, vmrseq is also applicable to the recently published sciEM-seq [40].

Nevertheless, we acknowledge that the evaluation through simulation studies potentially favors vmrseq, due to the fact that synthetic data were partly or fully generated according to assumptions of our model. Ideally, we would leverage alternative data-generative methods for a more unbiased method comparison and assessment. However, to the best of our knowledge, there are no existing models or functions in the literature that provide site-level single-cell bisulfite sequencing data. We recommend further evaluations using alternative methods if and when they become available.

Additionally, vmrseq is limited in that it relies on annotated datasets for parameter estimation (e.g., the transition probability and the beta priors in emission probability; see “Methods” and Additional File 1: Section S1.1). Due to lack of publicly available large-scale single-cell DNAme datasets at present, our training data in these analyses is concentrated in neuronal cell types. Applying vmrseq on tissues other than brain is essentially making the assumption that they share similar correlation and error structures to the neuronal subtypes. Results from Fong et al. [7] to some extent support the assumption of similarity between cell types in terms of between-CpG correlation distribution. Specifically, the authors used bulk whole-genome bilsulte sequencing data to study the autocorrelation of methylation level between CpGs with respect to genomic distances, and observed no significant difference in autocorrelations of methylation state across a range of cell types. We do not yet have evidence from single-cell data that shows consistency between cell types regarding the transition probability and beta-binomial distribution in emission probability. Future datasets generated on various cell types may be input to our package, vmrseq, for training relevant parameters and fill this gap.

Moreover, similar to scbs [23] and Smallwood et al. [9], vmrseq has limitations in that formal statistical significance testing (i.e., p-value quantification) is not available for region detection. This is due to the fact that CRs are already a set biased to high variability, precluding comparison of VMR statistics to a theoretical null. Further, the one-grouping HMM is not strictly a nested model of the two-grouping HMM hence likelihood ratio tests are not applicable. We do not see a straightfoward way to generate null distribution via either a theoretical or empirical approach. Detected VMRs can be ranked by the maximum likelihood increment from one- to two-grouping model; however, multiple testing corrections can not be applied thus false discovery rate is not strictly controlled. Extensions in this direction would provide a significant advance.

Last but not least, users of vmrseq should note that our method has only been experimented with and evaluated on autosomes. Sex chromosomes might have distinct methylation patterns from autosomes due to cellular processes such as the X-inactivation. Follow-up work is needed to examine whether model parameters for sex chromosomes shall differ from autosomes.

## Conclusions

We introduced vmrseq, a statistical approach with software implementation designed to discover variably methylated regions from single-cell bisulfite sequencing data; to the best of our knowledge this is the first such approach to do so at CpG site-level resolution. This method identifies biologically relevant regions at genome scale, overcoming extreme sparsity and technical noise inherent in single-cell technologies. Moreover, vmrseq is uniquely sensitive to the level of heterogeneity in input datasets. We extensively evaluated the performance of vmrseq using synthetic data and re-analyzed published large-scale studies of single-cell methylomes in mouse frontal cortex and gastrulation. In the simulation studies, our method outperformed previous approaches in a spectrum of accuracy metrics, demonstrating its utility in accurate region detection. In the case study of mouse cortex data, vmrseq was able to distinguish not only coarse cell classes but also detailed subtypes, outperforming the competing methods in terms of feature selection for clustering. In the case study of single cell multi-omic gastrulation, vmrseq identified heterogeneity associated with specific developmental stages and cell cycle states. Furthermore, our method elucidated intricate patterns of epigenetic regulation on gene expression that were not evident by looking only at promoters. In summary, vmrseq offers a reliable framework for pinpointing cell-to-cell DNA methylation heterogeneity, serving as a useful tool for future single-cell epigenomic research.

## Methods

### Overview of vmrseq

vmrseq takes processed and filtered whole-genome bisulfite sequencing data from heterogeneous individual-cell samples as input and performs a genome-wide scan for highly epigenetically variable regions. The methylation level of a CpG site in an individual cell is defined as methylated reads divided by total reads covering this site. Let *x_kc_*represent methylation level of CpG site *k* of cell *c*. The input to vmrseq is thus matrix *X* with element *x_kc_* on row *k* and column *c*. We assume binary methylation levels for each cell, i.e., *x_kc_* ∈ {0 (unmethylated), 1 (methylated)}. Missing is allowed in matrix *X* as vmrseq accommodates sparsity in scBS-seq data.

In practice, hemimethylation and technical error might occur and lead to intermediate methylation level between 0 and 1. However, only a small proportion of CpGs exhibit hemimethylation in most cells such as somatic mouse tissues [41]. In addition, the typical coverage level of scBS-seq data results in a large proportion of sites per cell (interquartile range 94.1–97.5% in datasets used in our case studies) only being observed by 1-2 read, precluding accurate identification of intermediate methylation in the first place. As a result, we remove the sites with an intermediate methylation level from every cell as a filtering step before input to vmrseq, which affects a very small proportion of sites per cell (interquartile range 0.33%–0.55%) in the datasets used in our case studies.

### Stage 1: construct candidate regions

#### Smoothing

vmrseq applies a kernel smoother to ‘relative’ methylation levels of individual cells to adjust for uneven coverage biases and borrow information from nearby sites. The relative methylation level of site *k* in cell *c* is defined as 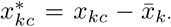, where *x̅_k_*_·_ is the average level across cells on that site and *c* ∈ {1, 2*, …, C*} with *C* being the total number of cells. Then for every cell, vmrseq runs a kernel smoother on 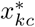, rendering smoothed values *x*^*_kc_*. We chose to smooth on the relative methylation levels instead of absolute ones to avoid introducing intermediate smoothed values unnecessarily in the cases where methylation state transitions between CpGs are consistent across the entire cell population. In all our studies, we adopted a box smoother to reduce computation time, and a bandwidth of 2,000 base pair (bp) was used since average correlation between CpG sites were observed to slowly decay as CpG-CpG distance grows and reach a plateau at around 2,000 bp (Additional File 1: Fig. S17). scbs [23] also employs relative methylation levels and kernel smoothing but in a different manner. In particular, scbs applies a smoother on the cell averages (i.e., *x̅_k_*_·_) and obtain the relative levels by subtracting individual-cell methylation by the smoothed average.

#### Defining candidate regions

Variance of methylation across cells is computed for each CpG site using the smoothed relative methylation. More formally, the variance is

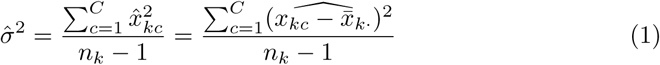

CRs are defined through identifying groups of at least 5 adjacent CpG sites whose variance is consistently greater than some threshold (Fig. 1b). This threshold is determined by taking the 1−*α* quantile value of an approximate null distribution of variance simulated from the beta priors of emission probability from our HMM (see Additional File 1: Section S1.2 for details). *α* is the designated significance level and its default is set to 0.05. The maximum distance between two adjacent CpG sites is set to 2,000 bp for the same reason of determining the default smoothing bandwidth. We also find that a value of 5 works well in practice as the minimum CpG counts in a CR and choose this for the default value. Please refer to Additional File 1: Section S2.4 for a sensitivity analysis on these hyperparameters. However, the specific application at hand can inform the choice of the minimum CpG count in a region. For example, a higher threshold (e.g., ≥ 50 CpGs) for capturing broad epigenetic changes in cancer studies [42], and a lower threshold (e.g., 5-10 CpGs) for detecting fine-scale regulatory changes in developmental studies [43].

### Stage 2: decode hidden Markov model

In the second stage, vmrseq detects VMRs inside each candidate region through decoding a hidden Markov model under the assumption of U and M groupings (Fig. 1c). HMMs are well-suited to model DNAme data, for they are in the form of sequences with the presence of local correlations. The HMM model used in vmrseq is motivated by the deconvolution method DXM [7]. Though DXM was designed for detecting heterogeneity in user-defined regions from bulk samples, we have discovered that a similar model is particularly relevant for detection of VMRs in single-cell data. We adopt the general structure of DXM, wherein the total observed methylated counts depends on the hidden states of CpG loci and association of adjacent CpG loci are modeled by transition probability between hidden states. But in contrast to DXM, which uses hidden states to directly infer cell subpopulations, we have proposed the assumption of U and M groupings and modeled the underlying methylation states of grouping(s) as hidden states. This is to accommodate the possibility that single-cell datasets might contain large number of cell subpopulations. To further adapt the method to single-cell technology, we adjust the model specification of transition and emission probabilities based on single-cell data characteristics (see later texts and Additional File 1: Section S1.1). We have also proposed an optimization step to estimate the prevalence parameter (detailed in later texts), instead of imposing a random choice a priori as in DXM.

We model the underlying methylation states of a grouping in a CR as a sequence of latent variables (i.e. hidden states in HMM) where each variable corresponds to a CpG site. Inference for each CR is made independently which enables the use of parallelization to lower computation time. We first describe the HMM for the one-grouping scenario (Additional File 1: Fig. S18a), as the two-grouping model (Additional File 1: Fig. S18b) is build upon the elements thereof.

#### One-grouping model specification

##### Hidden states

Denote the hidden states for site *k* with only one grouping as *s_k_* ∈ {0, 1} where 0 represents an unmethylated underlying state and 1 represents methylated. These states are considered latent because observed counts from data do not always align with the true methylation state due to both technical errors and inherent biological heterogeneity. The consecutive states are interdependent and are incorporated using transition probabilities to account for spatial correlations.

##### Transition probability

The transition probability from *s_k_*_−1_ to *s_k_* is denoted as *P* (*s_k_*|*s_k_*_−1_), which represents likelihood that the hidden state either changes or remains the same from one site to an adjacent site. The initial state probability is set to a non-informative uniform distribution, *P* (*s*_1_) ∝ 1. The transition probabilities are considered dependent on the methylation states and distance between two CpGs and are independent of the prevalence parameter. An empirical transition probability distribution used as default in vmrseq was trained from annotated single-cell bisulfite sequencing datasets previously published by Luo et al. [27] and Liu et al. [44] (see Additional File 1: Section S1.1.2 for training details, [45]).

##### Emission probability

For site *k*, denote its methylated cell count as 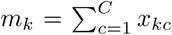 and total cell coverage as *n_k_* = |*ζ_k_*|, where *ζ_k_* is the set of cells with non-missing values at site *k* and | · | denote the cardinality of a set. The emission probability of observing *m_k_* methylated cell counts from *n_k_* total counts at site *k* is

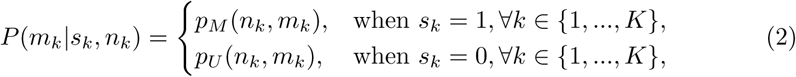

where *p_M_* (*n_k_, m_k_*) is a beta-binomial (BB) distribution modeling the *m_k_* out of *n_k_* cells *observed* as methylated given hidden state *s_k_* = 1. A separate zero-inflated betabinomial (ZIBB) distribution *p_U_* (*n_k_, m_k_*) models *m_k_* observed as unmethylated under the hidden state *s_k_* = 0. We use ZIBB for unmethylated states due to the fact that cell subtypes in labeled datasets [27, 35] were observed to exhibit unusually high proportion of zero counts hence not suitable for modeled as BB distribution. *p_M_* (·, ·) and *p_U_* (·, ·) together capture the measurement error that may occur during sequencing and inherent biological heterogeneity. Empirical prior distributions used as default in vmrseq was trained from annotated single-cell bisulfite sequencing datasets previously published by Luo et al. [27] and Liu et al. [44] (see Additional File 1: Section 1.1.3 for training details, [46, 47]).

##### Joint likelihood

For the *K* sites in a CR, denote the state sequence as **S** = (*s*_1_*, s*_2_*, …, s_K_*), methylated cell counts as **m** = (*m*_1_*, m*_2_*, …, m_K_*) and total cell count **n** = (*n*_1_*, n*_2_*, …, n_K_*). The likelihood for one grouping is consist of the transition and emission probability described above:

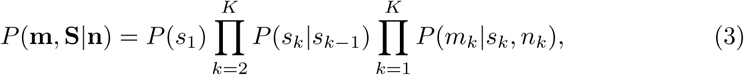

#### Two-grouping model specification

Recall the assumption of U and M grouping (Fig. 1d), where we assume that within a CR that contains a VMR, the cells can be partitioned into two groupings based on their estimated hidden states within the VMR (referred to as the U grouping and M grouping respectively). The two-grouping HMM is an extension of the one-grouping scenario, wherein each site possesses a bivariate hidden state and a prevalence parameter for depicting the proportion of cells in U and M groupings.

##### Hidden states

Denote the bi-variate hidden states for the two groupings as 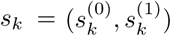 at site *k*, where 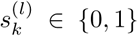 for *l* ∈ {0, 1}. Specifically, 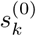 represents the underlying state of the U grouping at site *k*, and 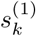 the underlying state of the M grouping. The state space is thus S = {(0, 0), (0, 1), (1, 1)}, where *s_k_* = (0, 1) indicates that the site *k* belongs to a VMR potentially (see later text for how a VMR is identified). *s_k_* = (0, 0) and *s_k_* = (1, 1) represents that site *k* is in the CR but not in a VMR. The combination (1, 0) is excluded from the state space here since we assume the M grouping must have a methylated hidden state at sites that are variably methylated. As a result of this constraint, the one-grouping HMM is not strictly nested to the two-grouping model.

##### Prevalence parameter

The prevalence parameter, i.e., the proportion of cells in the two groupings, is denoted as ***π*** = (*π*^(0)^*, π*^(1)^) with a constraint of *π*^(0)^ + *π*^(1)^ = 1.

##### Transition probability

Given a fixed between-CpG distance, the transition probability for two-grouping is extended from the *one-grouping transition probability*. Specifically,

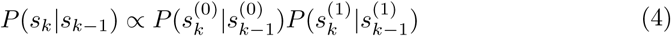

normalized by the constant 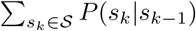 so that the sum equals 1. The initial state probability is also set to a non-informative uniform distribution, *P* (*s*_1_) ∝ 1.

##### Emission probability

For site *k* in a given candidate region, denote its methylated cell count as *m_k_* and total cell coverage as *n_k_* following the definitions in one-grouping case. The emission probability of observing *m_k_*methylated cell counts from *n_k_* total counts at site *k* is

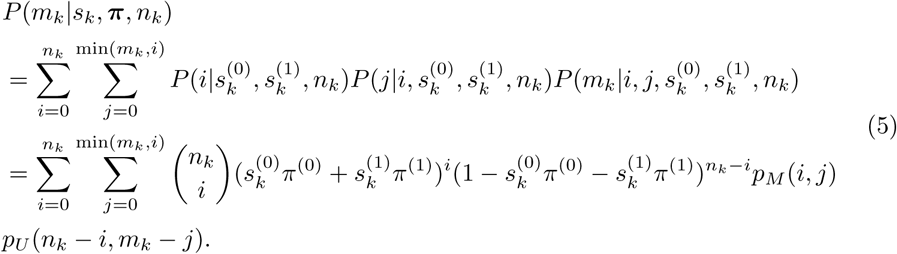

The first binomial term is for capturing the binomial sampling error where *i* cells came from the M grouping and *n_k_*− *i* come from the U grouping. Assuming cells are independently and identically distributed within a grouping, for site *k*, 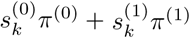 is the success rate in binomial distribution, representing the overall prevalence of cells with *underlying state of methylated*. Then, of the *i* cells, *j* may be *observed* as methylated, and is modeled by the BB distribution *p_M_* (*i, j*). Similarly, *p_U_* (*n_k_* − *i, m_k_* − *j*) is the ZIBB distribution representing the probability that *m_k_* − *j* of *n_k_* − *i* cells are observed as unmethylated. The BB and ZIBB models share the same structure and estimated parameters as in the one-grouping case.

### Joint likelihood

The joint likelihood can be written in the same manner as in the one-grouping case:

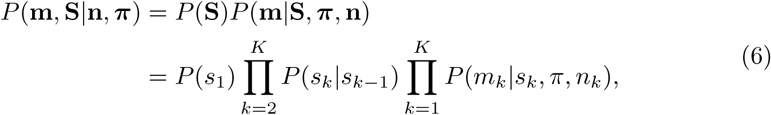

but with two-grouping representations of transition and emission probabilities instead.

#### Model optimization

In the one-grouping likelihood, the only parameters requiring estimation are the hidden states, as demonstrated in Eq. 3. Consequently, we can readily utilize the Viterbi algorithm [48] for HMM decoding. This algorithm is a dynamic programming technique that has traditionally been employed to obtain the maximum a posteriori probability estimate for the most probable sequence of hidden states. It is adopted here because of its excellent computational performance in terms of time complexity.

For optimizing the two-grouping likelihood, we repeatedly alternate between **S** and ***π*** to maximize the likelihood function for two groupings (Eq. 6) in a similar way as proposed by Rahman et al. [49]. We elaborate on these two phases of optimization in what follows.

##### 1. Initialization of π

Given the potential multimodality of the likelihood (Eq. 6), we recommend employing multiple initial values for the prevalence parameter. In practical applications, we have observed that the set {0.2, 0.5, 0.8} serves as an effective choice for initializing *π*^(0)^, and as a result, we have adopted it as the default setting in all experiments presented in this article.

##### 2. Optimizing S while π is fixed

Given the expected prevalence of two groupings, we solve for the most likely methylation state sequences in Eq. 6 by applying the Viterbi algorithm.

##### 3. Optimizing π while S is fixed

Maximizing the likelihood function over ***π*** is a constrained optimization problem with ***π*** as a probability vector, where *π*^(0)^*, π*^(1)^ are non-negative and *π*^(0)^ + *π*^(1)^ = 1. To solve this problem, we use the exponentiated gradient (EG) algorithm. More formally, the optimization problem with respect to ***π*** is as follows:

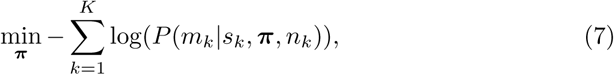

in which the objective function is proportional to the logarithm of likelihood function in Eq. 6. Then, the EG updates are

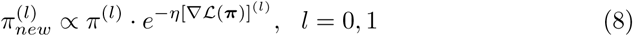

where *η* is the learning rate. After updating each component of the latent vector ***π***, the values are normalized so that they sum to one. The derivatives in EG updates are provided in Additional File 1: Section S1.3.

##### 4. Termination

For each initialization of *π*^(0)^ (step 1), continue iterating through steps 2 and 3 until convergence is achieved. The optimization process is considered complete when both the objective function and *π*^(0)^ have converged. Select the parameter estimates with the highest maximum likelihood among the fitted models generated from all initializations.

When sparsity level in the input dataset is held constant, the observed total cell count *n_k_*should exhibit linear scaling with the total number of cells, *C*. Consequently, the time complexity for optimizing the one-grouping Hidden Markov Model (HMM) is *O*(*KC*^2^), while for the two-grouping HMM, it becomes *O*(*IKC*^2^), where *I* represents the total number of iterations.

#### Identifying and ranking variably methylated regions

For a candidate region, if the one-grouping HMM achieves a higher maximum likelihood than the two-grouping HMM, this CR is not considered to contain VMR. Otherwise, a VMR is defined for regions with a minimum number of CpG sites exhibiting hidden state 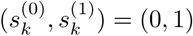. Same as in candidate region construction, we used 5 as the minimum number for all experiments in this study and it can be customized by the user based on the specific biological question at hand. Should multiple VMRs be detected in one CR, we merge those VMRs into one to be concordant with the assumption of U and M groupings. The VMRs can be ranked by the increment in log-likelihood of two-grouping model compared to one-grouping with a higher increment indicating a higher rank.

#### Annotated datasets used for empirical parameter estimation

The single-cell bisulfite sequencing datasets published by Luo et al. [27] and Liu et al. [44] were used for training the empirical parameters in transition and emission probability distributions in our model. Originally, Luo et al. [27] profiled over 6,000 single neuronal nuclei methylomes using the snmC-seq protocol, identifying 16 mouse and 21 human neuronal subtypes in the frontal cortex using gene body non-CG methylation depletion in neuronal marker genes. We also re-analyzed this dataset in the first case study. Similarly, Liu et al. [44] profiled more than 100,000 nuclei with snmC-seq2, including both neurons and non-neuronal cells, from 45 distinct regions of the mouse brain. They identified 161 cell subtypes with high consistency across different replicates, which are characterized by unique spatial distributions and projection targets, leveraging both CG and non-CG methylation patterns. Cells from each annotated subtype were merged into an individual pseudo-bulk sample and these samples are considered homogeneous during empirical parameter estimation. For specific details of data processing and parameter training, please refer to Additional File 1: Section S1.1.

#### Computational scalability of vmrseq

We conducted a scalability analysis using synthetic data to evaluate the running times of various methods across different settings (Additional File 1: Fig. S19). Due to the second-order computational complexity with respect to the number of cells, the running time of vmrseq exhibits a quadratic increase as cell count grows, while vmrseq CRs, scMET and scbs present a linear increase. Therefore, when dealing with datasets consisting of a large number of cells, we recommend running only the first stage of the methodology for users who need a preliminary check or prioritize speed over high precision in region detection and the ranking of regions. This approach allows for a faster initial analysis before applying the full methodology.

### Method implementations and parameter settings

The alternative methods that were compared with vmrseq were exclusively applied in the context of a genome-wide search for VMRs, regardless of other functions they might possess. All analyses were carried out using R version 4.2.0 [50] except for the application of the scbs which is a stand-alone command-line tool developed from Python. We used default parameters as suggested in original publications of the evaluated methods in all our experiments. For RRA estimation, the sets of thresholds were tailored to each method to balance the trade-off between computational time and a PR-curve with steady and continuous progression. The sites with an intermediate methylation level were excluded for every cell before input to any of the methods. In both datasets used in the case studies, 0.44% of sites were removed on average from each cell (interquartile range 0.33–0.56%). Method-specific implementations and parameter settings are described as follows.

#### vmrseq

The `vmrseq` package (https://github.com/nshen7/vmrseq; version 0.1.0) was used for analyses in this article with default parameters (smoothing bandwidth = 2000 bp; variance quantile *α* = 0.05; minimum number of CpGs in CR = 5; minimum number of CpGs in VMR = 5). For computing RRA of precision-recall curve, *α* ∈ {.001, .002, …, .005, .01, .02, …, .1, .12, .15, .2, .3, .4} were used. *α*’s that are greater than 0.4 were not included because large values of *α* might not effectively define candidate regions that align with our assumption of U and M groupings. For all experiments, we removed CpGs sites with less than 3 covered cells prior to applying vmrseq.

#### scbs

scbs [23] first computes cell-to-cell variances of smoothed mean methylation levels in fixed-size sliding windows, then takes a user-defined percentage of top-ranked windows in terms of the variance and merges them whenever overlapping happens. The non-overlapping regions after merging are identified as VMRs and are re-ranked based on across-cell variance in methylation. The scbs package (https://github.com/LKremer/scbs; version 0.4.0) was used for implementation. The default 2000-bp window size, 10-bp step size were used for the all experiments in this article. Default 2% variance threshold (i.e. proportion of top-ranked windows) were used for all experiments except that for computing RRA of precision-recall curve, variance cutoffs were {.0001*, .*0005*, .*001*, .*005*, .*01*, .*015*, .*02*, .*025*, .*03*, .*04*, …, .*09*, .*1*, .*2*, …, .*9*, .*99}. No additional pre-processing steps were used for scbs. On a side note, the scbs method has been updated and renamed as MethSCAn in its latest version [51], following the completion of our experiments.

#### Smallwood

The statistical method used in Smallwood et al. [9] models the aggregated methylation counts from fixed-size sliding windows with a binomial distribution. Regions are ranked by the confidence lower bound of their maximum likelihood estimator of the cell-to-cell methylation variance and an arbitrary number of top-ranked regions are defined as variably methylated. Selected windows that overlapped are merged into non-overlapping regions. Relying on the description provided in Smallwood et al. [9], we created our own implementation as we were unable to find a readily available software package to use. As proposed in their study, 3,000-bp window size with 600-bp step size were applied for the studies in this article. However, we performed additional experiments using the same 2,000-bp window size as vmrseq and scbs to assess the impact of varying window size on method performance (see Additional File 1: Section S2.1 for a detailed discussion). The article originally used top 300 windows, which we deemed a too small number for largescale datasets, thus top 2% windows were used instead as default (in light of the threshold proposed by scbs). For computing RRA of precision-recall curve, variance cutoffs {.0001*, .*0005*, .*001*, .*005*, .*01*, .*015*, .*02*, .*025*, .*03*, .*04*, …, .*09*, .*1*, .*2*, .*3} were used. No additional pre-processing steps were used.

#### scMET

scMET [26] leverages the concept of ‘residual overdispersion’ that supposedly removes the confounding of variance by mean methylation and models the aggregated counts in pre-defined windows with a hierarchical Bayesian model. The `scMET` package (https://www.bioconductor.org/packages/release/bioc/html/scMET.html; version 0.99.11) was used for implementation. As proposed in their study of genome-wide sliding windows, 20,000-bp window size with 20,000-bp step size were used for the studies in this article, along with default parameters in `scmet` and `scmet_hvf` function of `scMET` package. In particular, the default threshold used for selecting highly variable windows is expected false discovery rate (EFDR) = 0.1; and for computing RRA of precision-recall curve, EFDR of values {.01*, .*02*, .*05*, .*1*, .*2*, …, .*9, 0.99} were used. For all experiments, we followed the recommended processing steps from [26] and removed windows with less than 3 covered sites for each cell and features that did not have CpG coverage in at least 5 cells.

### Statistical analysis

#### Ratio of relevant areas (RRA)

Because the identification of VMRs requires an increase of likelihood from a one-grouping to a two-grouping model in vmrseq, the precision is not guaranteed to reach 0 by lowering the variance threshold; hence a precision-recall curve with precision ranging from 0 to 1 inclusive can not be drawn for VMRs detected by vmrseq. Therefore, instead of using area under precision-recall curve as the criterion to comprehensively compare accuracy of the methods, we applied RRA, a metric that restricts the evaluation of area under curve to a region of interest. Specifically, we set the region of interest as 0.8 ≤ precision ≤ 1 for the experiments with synthetic data to assess performance under low-false-discovery-rate (i.e., high-precision) conditions while accommodating the aforementioned circumstance of vmrseq.

#### Cell-to-cell dissimilarity matrix

Given a genomic region, for each cell, regional average methylation was computed by taking the mean observed methylation of all CpGs in the region. The cell-to-cell dissimilarity matrix was composed of pairwise Manhattan distances (i.e., *L*_1_ distance) between the regional average methylation vectors of every possible pair of cells. For each cell-cell pair, only regions observed in both cells were included in calculation of distance. The implementation was performed using daisy function from `cluster` package in R (version 2.1.3).

#### Dimension-reduction visualization

For each method, UMAP [28] was applied directly to the dissimilarity matrices computed from detected regions to obtain embedded coordinates in 2-dimensional space. The implementation was performed using the `umap` function from the `uwot` package in R (version 0.1.14) with default parameters. The same random seed was used for all methods in the studies.

#### Nearest neighbor count score

For each cell *c*, let *γ_c_* represent the count of its *g* nearest neighbors (determined by the dissimilarity matrix) that possesses the same label as *c*. A cell was deemed “well-assigned” if *γ_c_ > θg*, where we set *g* = 100 and *θ* = 0.7 in the case study. The nearest neighbor count score was defined as the fraction of cells considered well-assigned. This score assesses the capability of each method to distinguish between cell types in the detected regions. Each score ranges from 0 to 1, with higher scores indicating superior separation. Other parameter settings of *g* and *θ* rendered very similar results on the mouse multi-omic dataset (Additional File 1: Fig. S20).

### Generating synthetic datasets

To ensure that the simulated sets closely match the characteristics of the observed experimental data, we replicated chromosome 1 of subtype ‘IT-L23 Cux1’ from an scBS-seq mouse brain atlas [44] in terms of genomic coordinates and sparsity distribution. Cells of only one subtypes were used to ensure no known heterogeneity was present in the dataset. We stratified the total 6,550 cells from this cell subpopulation into three subsets based on their sparsity levels, where the level of sparsity of a cell was defined as the percentage of CpG sites not covered by any read. The top and bottom 2% of cells were excluded to remove potential outliers. 2,096 cells remaining in each subset and average sparsity levels from the three subsets are 94.9% (high), 92.8% (medium), 90.4% (low) respectively. We randomly subsampled N = 200, 500, 1000, 2000 cells from each of the three subsets, resulting in 12 sets of cells for synthetic purposes. Additionally, a range of values for number of subpopulations (denoted as *ρ*; *ρ* ∈ {2, 3, 4, 5, 8, 12, 20}) were simulated for each of the 12 sets.

Assuming that these sets are homogeneous and contain no VMRs, we inserted 2000 simulated VMRs into each set. VMR positions were determined by sampling clusters of between 5 and 500 CpGs with a maximum gap between any two adjacent CpGs of 500 base pairs and maximum cluster width of 10,000 bp. Adjacent VMRs with ≤ 2 CpGs in between are merged. Using a simulation procedure similar to Korthauer et al. [12], we prioritized the selection of clusters with mean methylation averaged across all CpGs near 0.5. Mean methylation of a CpG is calculated as the number of methylated cells divided by the total covered cells of a site. This approach was adopted to more accurately capture observed biological variability, as sample-to-sample variability tends to be high at intermediate methylation levels [52, 53]. This preference was enforced using probability weights *w_r_, r* = 1*, …, R* when sampling over the possible *R* CpG clusters, where 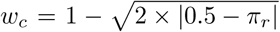 and *π_r_* is the mean methylation over all sites in cluster *r*.

According to the assumption of U and M groupings, all sites within VMRs had two groupings and fixed hidden states (*s*^(0)^*, s*^(1)^) = (0, 1). We sampled the prevalence parameter based on *ρ*. Specifically, for each VMR, we took a random number of subpopulations (denoted as *ρ_r_*; 0 *< ρ_r_ < ρ*) to be in M grouping and others in U grouping. Then the prevalence of U grouping for this VMR, 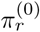, followed the distribution:

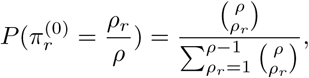

and 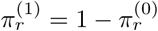 accordingly.

For each VMR, we then randomly assigned a proportion of cells to the U and M groupings according to the sampled prevalence parameter. For each CpG site in U and M grouping, the methylated cell count (conditional on observed total cell coverage) was drawn from the beta-binomial components in the emission probability of HMM in vmrseq stage 2, *p_U_* (·, ·) and *p_M_* (·, ·), respectively. At last, those methylated cell quotas were randomly assigned to the cells for each site, excluding the cells missing from this site due to zero-read coverage.

For the second set of experiments wherein methylation levels of the whole chromosome are entirely simulated from the HMM model, VMRs were generated in the same manner. Sites outside the VMRs were assumed to only contain a single grouping. The hidden states of these sites were sampled from the transition probability based on observed between-CpG distances. Then the cell methylation values were simulated from the emission probability distribution based on the generated hidden states.

### Mouse frontal cortex dataset

#### Data processing

CpG methylation read counts per cell following alignment to the mm10 mouse genome are available from the Gene Expression Omnibus repository under accession number GSE97179. We included the same set of cells as used in Kapourani et al. [26]. Details on quality control and data pre-processing on raw reads can be found in Luo et al. [27]. See Additional File 2: Table S2 for sample metadata of the included cells. Only autosomes were included in this study.

#### Annotations of detected regions with genic contexts and CpG region types

We annotated detected regions from each method by overlapping with 11 types of gene contexts and 4 types of CpG region types. The gene context annotations were obtained using the “genes 1to5kb”, “genes promoters”, “genes cds”, “genes 5UTRs”, “genes exons”, “genes firstexons”, “genes introns”, “genes intronexonboundaries”, “genes exonintronboundaries”, “genes 3UTRs” and “genes intergenic” annotations of genome “mm10” from version 1.22.0 of the `annotatr` package in Bioconductor [54]. The CpG region type annotations (including islands, shores, shelves and inter-island areas) were obtained using “cpgs” annotations from the same package. Randomized regions were obtained by employing function `randomize regions` from the same package.

### Multi-omics gastrulation dataset

#### Data processing

CpG methylation read counts per cell following alignment to the mm10 mouse genome were downloaded from ftp://ftp.ebi.ac.uk/pub/databases/scnmt_gastrulation/scnmt_gastrulation.tar.gz. Raw sequencing reads are available from the Gene Expression Omnibus repository under accession number GSE121708. Details on the quality control and data processing can be found in Argelaguet et al. [32]. In total 939 cells remained after taking the subset of cells with both QC-passed methylomic and transcriptomic profiles available. See Additional File 2: Table S3 for sample metadata of the remaining cells. As with the reanalysis on [27] dataset, only autosomes were included in this study. Genes with fewer than 5 cells of coverage were excluded, leaving a total of 17,568 genes for downstream analysis.

#### Annotation of cell cycle phases

The cell cycle phases of individual cells were annotated using the `CellCycleScoring` function from R package `Seurat` (version 4.3.0), following the steps in vignette https://satijalab.org/seurat/articles/cell_cycle_vignette.html. The annotation was performed based on the mouse ortholog genes of the human cell cycle canonical markers used in this vignette. The `gorth` function from the R package `gprofiler2` (version 0.2.2) was used for the orthology search. See Additional File 1: Fig. S15a for distribution of S and G2M scores on the annotated cells.

#### Selection of highly variable genes

Gene expression of individual cells were first normalized with variance stabilizing transformation via vst function from Bioconductor package `sctransform`. Genes in the top 10% ranked by variance of normalized gene expression across cells were selected in the case study. As a sanity check, we performed gene ontology enrichment analysis using the list of selected highly variable genes. As expected, most gene pathways associated with these genes are relevant to embryonic organ development and gastrulation (Additional File 1: Fig. S16a).

#### Linking VMRs to genes

Only VMRs with any boundary located within a distance of 1,000 bp to a gene (ranging from transcription start site to transcription end site) was considered linked to that gene. A promoter was defined as 2, 000 bp upstream to the transcription start site of the corresponding gene. A VMR was assigned to category “overlapping with promoter” if any of its boundary was in a promoter. A VMR was assigned to “overlapping with gene body” if any of its boundary was in a gene body but none of the boundaries was in promoter. At last, a VMR was assigned to “outside the gene but within 1000-bp distance” if none of its boundary was in promoter or gene body.

#### Correlation of regional average methylation and gene expression

For both VMRs and promoters, the Spearman correlation of regional average methylation and normalized expression of linked gene was computed using cells with observations of both methylation level and gene expression. VMRs and promoters with less than 10 cells covered were removed from the analysis to ensure a robust evaluation of correlation. Additional File 2: Table S4 contains information of all gene-VMR pairs along with the computed correlations.

#### Gene ontology enrichment analysis

All enrichment analyses in the experiments were performed using the enrichGO function from Bioconductor package `clusterProfiler` (version 4.4.4) with organism annotation database `org.Mm.eg.db` (version 3.15.0).

## Declarations

## Ethics approval and consent to participate

Not applicable.

## Consent for publication

Not applicable.

## Availability of data and materials

All datasets used in this article are publicly available. The mouse cortex dataset from Luo et al. [27] is available under GEO accession number GSE97179 [55]. The parsed scNMT-seq gastrulation dataset [32] was downloaded from ftp://ftp.ebi.ac.uk/pub/databases/scnmt_gastrulation/scnmt_gastrulation.tar.gz. Raw files are available under GEO accession number GSE121708 [56]. The mouse brain dataset from Liu et al. [44] is available under GEO accession number GSE132489 [57]. The vmrseq framework is publicly available as R software released under the MIT license (see https://github.com/nshen7/vmrseq [58]). A snapshot of the software used for generating results in this manuscript can be found at https://doi.org/10.5281/zenodo.11556577 [59]. The code used to process and analyze the data is available at https://github.com/nshen7/vmrseq-experiments [60] under the MIT license, and a snapshot is available at https://doi.org/10.5281/zenodo.14226597 [61].

## Authors’ contributions

KK conceptualized the problem. NS and KK developed the methodology. NS implemented the method and wrote the R package. NS conducted all experiments and drafted the manuscript. KK and NS contributed to reviewing and editing the manuscript. All authors have approved the final manuscript.

## Acknowledgements

This research was supported in part through the computational resources and services provided by Advanced Research Computing at the University of British Columbia.

## Acronyms

BB: beta-binomial. 17, 18
bp: base pair. 15, 21, 22, 25
BS-seq: bisulfite sequencing. 1
CR: candidate region. 3–5, 7, 13, 15, 16, 20, 21
DNAme: DNA methylation. 1–3, 10–13, 16
EG: exponentiated gradient. 19
HMM: hidden Markov model. 4, 5, 7, 13, 15–17, 24
PCA: principal component analysis. 9
RRA: ratio of relative areas. 5, 7, 21, 22
VMR: variably methylated region. 2–7, 9–13, 16, 20–25
ZIBB: zero-inflated beta-binomial. 17, 18

