## Additional File 1 for "vmrseq: probabilistic modeling of single-cell methylation heterogeneity"

### Supplementary material for **vmrseq: probabilistic modeling of single-cell methylation heterogeneity**

### S1 Supplementary methods

#### S1.1 Empirical parameter estimation in HMM

##### S1.1.1 Processing the annotated datasets

We make use of labeled single-cell methylation data from Luo et al. [1] and Liu et al. [2] for the estimation of transition probability and beta priors of emission probability. Luo et al. [1] profiled in total $> 6,000$ methylomes from single neuronal nuclei using snmC-seq and annotated 16 mouse and 21 human neuronal subtypes in the frontal cortex based on gene body non-CG methylation depletion in neuronal marker genes. Liu et al. [2] profiled $> 100,000$ nuclei with snmC-seq2 (including neurons and non-neuronal cells) from 45 dissected regions of the mouse brain. 161 cell subtypes with distinct spatial locations and projection targets were annotated based on CG and non-CG methylation, where all subtypes are highly conserved across replicates.

The empirical parameters are trained on subtype-specific pseudo-bulk data aggregated from Luo et al. [1] and Liu et al. [2]. A substantial proportion of subtypes in these datasets contain a small number of cells. Specifically, 82 out of total 161 subtypes contain less than 300 cells. Using all the subtypes might cause an imbalanced model that disproportionately emphasizes the smaller subtypes. Therefore, to ensure a balanced distribution of cell count, we divided the subtypes into groups with fixed range of cell counts (e.g., (0, 100], (101, 200], ...) and subsampled the subtypes by randomly selecting at most 3 within each group. As a result, cell counts of the remaining subtypes are approximately uniformly distributed in the range from 46 to 6,550. In a similar way, we included a subset of subtypes from Luo et al. [1] data in parameter training. Please see Additional File 2: Table S5 for the list of subtypes selected to train the parameters.

Notation in this section follows the "Methods" section of the main text. Given a subtype $t$, we sum up the single-cell binary methylation levels $x_{kc}$ to obtain methylated cell count $m_k^t = \sum_{c \in C_t} x_{kc}$ and total cell coverage $n_k^t = |\zeta_k^t|$ respectively for each CpG site $k$, where $\zeta_k^t$ is the set of cells with non-missing values at site $k$ for subtype $t$, and $|\cdot|$ denote the cardinality of a set. For each cell subtype, CpG sites with total cell coverage $n_k^t < 5$ to reduce impact of uncertainty due to technical noise.

The underlying methylation state of site $k$ in subtype $t$ is defined as

$$\tilde{s}_k^t = \begin{cases} 1, & m_k^t/n_k^t > 0.5 \\ 0, & m_k^t/n_k^t < 0.5 \\ \tilde{s}_{k-1}^t, & m_k^t/n_k^t = 0.5 \end{cases}$$

##### S1.1.2 Estimation of transition probability

The transition probability quantifies the likelihood of a change in underlying methylation states from one CpG to another within a grouping. Generally, a decreasing correlation between methylation levels of CpGs is expected as the genomic distance between the CpG pair increases (Fig. S17). Hence, we model the transition probability as a function of between-CpG distance and obtain the estimates by applying local polynomial regression. We first describe transition probability estimation for one grouping, since the transition probability in the two-grouping model is derived from it.

Genomic distance $d_{(j_1,j_2)}$ between two CpG sites $j_1$ and $j_2$ on the same chromosome (where $j_1 < j_2$) is measured as the difference of their genomic coordinates in base pairs. We take CpG pairs that are not farther apart than 10 sites, i.e., $j_2 - j_1 \leq 10$ and denote $\eta_d$ as the set of all applicable $(j_1, j_2)$ with respect to distance $d$.

For example, for subtype $t$ we may estimate the probability of transitioning from an unmethylated state (i.e., $s_{k-1} = 0$) to methylated (i.e., $s_k = 1$) as a function of distance $d$:

$$P_d^t(s_k = 1 | s_{k-1} = 0) = a_{01}^t(d) = \frac{\sum_{(j_1,j_2)\in\eta_d} \mathbf{1}\{\tilde{s}_{j_1}^t = 0 \text{ and } \tilde{s}_{j_2}^t = 1\}}{\sum_{(j_1,j_2)\in\eta_d} \mathbf{1}\{\tilde{s}_{j_1}^t = 0\}}.$$

Then we fit a LOESS [3] smoothing curve on the average probability across subtypes $a_{01}(d) = \frac{1}{T}\sum_{t=1}^{T} a_{01}^t(d)$ with respect to $d$, with inverse variance weights $w_{01}(d) = (\frac{1}{T-1}\sum_{t=1}^{T}(a_{01}^t(d) - a_{01}(d))^2)^{-1}$ using R function `loess` in package `stats` (version 4.2.0), where the locally fitted values from LOESS are denoted as $\hat{a}_{01}(d)$ with respect to distance $d$. The degree of polynomial and span was set as 2 and 0.03 respectively. Distance $d$ was set on the log scale during the curve fitting to improve the modeling of the relationship between transition probability and between-CpG distance, particularly when the distance is short (Fig. S14). Fitted values from LOESS across subtypes are considered estimated transition probabilities from state 0 to state 1, i.e., $P_d(s_k = 1|s_{k-1} = 0) = \hat{a}_{01}(d)$ with respect to $d$.

Similarly we may also estimate $P_d(s_k = 0|s_{k-1} = 0) = \hat{a}_{00}(d)$, $P_d(s_k = 0|s_{k-1} = 1) = \hat{a}_{10}(d)$ and $P_d(s_k = 1|s_{k-1} = 1) = \hat{a}_{11}(d)$. Transitions with a distance greater than 2,000 bp are considered fixed and identical to distribution at 2,000 bp because the estimated probability usually reaches a plateau as distance exceeds 2,000 bp (Fig. S14; Additional File 2: Table S6). In summary, we estimated empirical transition probabilities, $P_d(s_k = 0|s_{k-1} = 0)$, $P_d(s_k = 0|s_{k-1} = 1)$, $P_d(s_k = 1|s_{k-1} = 0)$ and $P_d(s_k = 1|s_{k-1} = 1)$ with respect to $d = 2, 3, 4, ..., 2000$ bp via LOESS using the pseudo-bulk samples aggregated from the annotated single-cell datasets. These empirical probabilities were used as default throughout this study.

##### S1.1.3 Estimation of beta priors in emission probability

***Estimation of*** $p_U(\cdot, \cdot)$

To obtain $p_U(\cdot, \cdot)$, we train a zero-inflated beta-binomial (ZIBB) generalized linear model [4] on $m_k^t$ of sites with underlying unmethylated state with respect to logarithm of $z^t$, where $z^t$ is the median coverage across all CpG sites in subtype $t$. In other words, $z^t = \text{median}\{n_k^t : k \in \kappa\}$, where $\kappa$ represents the complete set of CpG sites in the genome. The intuition behind including the covariate $z^t$ is that we think a higher amount of inherent biological heterogeneity shall be allowed in a larger cell grouping. We consider median coverage to be a more suitable representation of the grouping's magnitude than the total number of cells because coverage takes into account both the cell count and the sparsity level. More formally, the ZIBB regression model has the following formulation.

We first introduce the ZIBB distribution. Let $Y = 0$ with probability $\nu$ and $Y \sim \text{BB}(n, \mu, \sigma)$ with probability $1 - \nu$, where $0 < \mu < 1$ and $\sigma > 0$ are the mean and precision parameter in the beta prior. Then $Y$ follows the ZIBB distribution, denoted by $\text{ZIBB}(n, \mu, \sigma, \nu)$, given by:

$$p_Y(m|n, \mu, \sigma, \nu) = \begin{cases} \nu + (1-\nu)p_{Y'}(0|n, \mu, \sigma), & \text{if } m = 0 \\ (1-\nu)p_{Y'}(m|n, \mu, \sigma), & \text{if } m = 1, 2, 3, ..., \end{cases}$$

where $Y' \sim \text{BB}(n, \mu, \sigma)$.

We assume that $m_k^t \sim \text{ZIBB}(n_k^t, \mu_k^t, \sigma_k^t, \nu_k^t)$ with a linear effect of a cubic spline of median coverage $z^t$. where $\mu$ and $\nu$ are logit link-transformed and $\sigma$ are log link-transformed. These modeling choices result in the following generalized linear model:

$$\begin{aligned} \text{logit}(\nu_k^t) &= \beta_{\nu,0}^U + \beta_{\nu,1}^U S(\log(z^t)) \\ \text{logit}(\mu_k^t) &= \beta_{\mu,0}^U + \beta_{\mu,1}^U S(\log(z^t)) \\ \log(\sigma_k^t) &= \beta_{\sigma,0}^U + \beta_{\sigma,1}^U S(\log(z^t)) \end{aligned} \tag{S1}$$

where $S(\cdot)$ represents cubic splines. We used $(m_k^t, n_k^t, z^t)$ for training, where $t \in \{1, ..., T\}, k \in \{1, ..., K_{\text{sub}}\}$. We randomly subsampled $K_{\text{sub}} = 10,000$ sites with an underlying unmethylated state (i.e., $\tilde{s}_k^t = 0$) from each subtype $t$ to alleviate the computational burden of model optimization during training. In summary, the regression intercepts $\{\beta_{\nu,0}^U, \beta_{\mu,0}^U, \beta_{\sigma,0}^U\}$, slopes $\{\beta_{\nu,1}^U, \beta_{\mu,1}^U, \beta_{\sigma,1}^U\}$ and cubic spline function $S(\cdot)$ were estimated via maximum likelihood method.

With the trained ZIBB regression model, we may predict the beta prior parameters for an input dataset, $\nu, \mu$ and $\sigma$, based on its median cell coverage $z$:

$$\begin{aligned} \text{logit}(\nu_k^t) &= \hat{\beta}_{\nu,0}^U + \hat{\beta}_{\nu,1}^U \hat{S}(\log(z^t)) \\ \text{logit}(\mu_k^t) &= \hat{\beta}_{\mu,0}^U + \hat{\beta}_{\mu,1}^U \hat{S}(\log(z^t)) \\ \log(\sigma_k^t) &= \hat{\beta}_{\sigma,0}^U + \hat{\beta}_{\sigma,1}^U \hat{S}(\log(z^t)) \end{aligned}$$

See Fig. S21a-c and Additional File 2: Table S7 for estimated beta prior parameters with respect to $\log(z^t)$. The range of $z$ is limited to $[0, 400]$ due to the size of the training data. Since the estimated parameters seem to level off as $z$ increases (Fig. S21a-c), we use the same set of parameters from $z = 400$ for the cases $z > 400$.

###### *Estimation of* $p_M(\cdot, \cdot)$

Originally the same modeling choice was tried for $p_M(\cdot, \cdot)$. However, based on model diagnostics, $z^t$ does not seem to play a significant role in the distributional parameters of $p_M(\cdot, \cdot)$ among sites with underlying methylated state (i.e., $\tilde{s}_k^t = 1$). Therefore, we model $p_M(\cdot, \cdot)$ as a intercept-only BB regression model, i.e.,

$$\begin{aligned} \text{logit}(\mu_k^t) &= \beta_{\mu,0}^M \\ \log(\sigma_k^t) &= \beta_{\sigma,0}^M \end{aligned} \tag{S2}$$

which is trained on $(m_k^t, n_k^t, z^t)$, where $t \in \{1, ..., T\}, k \in \{1, ..., K_{\text{sub}}\}$. The parameters $\beta_{\mu,0}^M$ and $\beta_{\sigma,0}^M$ were estimated via maximum likelihood. Similarly, $K_{\text{sub}} = 10,000$ sites with $\tilde{s}_k^t = 1$ were randomly subsampled from each subtype for training. See Fig. S21d-e and Additional File 2: Table S8 for estimated beta prior parameters with respect to $\log(z^t)$.

###### *Implementation*

R packages `gamlss` (version 5.4-3) and `gamlss.dist` (version 6.0-3) were used for implementation for optimizing both $p_U(\cdot, \cdot)$ and $p_M(\cdot, \cdot)$. The default Rigby and Stasinopoulos algorithm [5] was used for maximum likelihood optimization.

##### S1.1.4 Sensitivity analysis on empirical parameter estimation

In estimating transition probabilities and beta priors in emission probabilities, we utilized the datasets from Luo et al. [1] and Liu et al. [2] to maximize the training set size and enhance generalizability. However, since the Luo et al. [1] dataset was also employed for benchmarking the performance of vmrseq and alternative methods, we conducted a sensitivity analysis to evaluate the empirical parameter estimation's robustness against the datasets used for training. Specifically, we examined the difference between the sets of estimated parameters before and after the Luo et al. [1] dataset being excluded from training.

Fig. S22 and S23 respectively display the estimated transition probability and beta priors in emission probability before and after the exclusion of the Luo et al. [1] data. The transition probability estimates derived solely from the Liu et al. [2] dataset are particularly close to the original estimate obtained using both datasets. The training of beta prior parameters in emission probability also yields estimates that are in close proximity to the original estimates. This demonstrates that the estimation process is fairly robust to the exclusion of Luo et al. [1] data during training.

#### S1.2 Variance threshold determination for candidate region construction

Under the assumption of U and M grouping, the null condition presumes that all CpG sites in the genome have only one grouping. In other words, a site should have underlying state of either methylated or unmethylated. With this premise, we use Monte Carlo simulation to generate an empirical null distribution of cell-to-cell variance given the coverage level of an input dataset. Specifically, we first sample values of variance from underlyingly methylated and unmethylated sites separately, and then concatenate them to form the complete empirical distribution. Subsequently, the threshold on variance is determined by taking the quantile $1-\alpha$ in this empirical distribution, where $\alpha$ is a small value bewteen 0 and 1 (default at $\alpha = 0.05$). Thus, exceeding this quantile implies an *unusually* high cell-to-cell variance. More formally, the null distribution of variance is generated through the following steps. Denote the methylated fraction of cells for a site $k$ as $\bar{x}_{k\cdot} = \frac{1}{C}\sum_{c=1}^{C} x_{kc}$, where $x_{kc}$ is the binary methylation level of site $k$ and cell $c$.

1. Use sites with relatively extreme methylated fractions to estimate proportions of methylated and unmethylated hidden states, $(\phi_0, \phi_1)$, for the genome under null condition:

$$\phi_0 = \frac{\sum_{k\in\kappa} \mathbf{1}\{\bar{x}_{k\cdot} < 0.4\}}{\sum_{k\in\kappa} \mathbf{1}\{\bar{x}_{k\cdot} < 0.4\} + \sum_{k\in\kappa} \mathbf{1}\{\bar{x}_{k\cdot} > 0.6\}}$$

   and $\phi_1 = 1-\phi_0$, where $\kappa$ represents the complete set of CpG sites in the genome.
2. Compute the logarithm of median across-cell coverage $z$ of user input dataset. Obtain beta prior parameters in $p_U(.,.)$ and $p_M(.,.)$ based on the trained emission probability distribution and $z$, denoted as $(\nu_U, \mu_U, \sigma_U)$ and $(\mu_M, \sigma_M)$ respectively (see section S1.1.3).
3. Let $V$ be number of sampled values (default to $V = 100,000$). Sample $\phi_0 V$ times from zero-inflate beta distribution with parameter $(\nu_U, \mu_U, \sigma_U)$ and $\phi_1 V$ times from beta distribution with parameter $(\mu_M, \sigma_M)$. Concatenate these two set of sampled values, denoted as $\{p_1, ..., p_V\}$.
4. Considering methylation level of any CpG site $x_{k\cdot}$ as a binary variable following the Bernoulli distribution, i.e., $x_{k\cdot} \sim \text{Bernoulli}(p)$, the variance of $x_{k\cdot}$ shall be

$p(1-p)$. Hence, the null distribution of variance follows the empirical distribution of $\{p_1(1-p_1), ..., p_V(1-p_V)\}$; and the threshold on variance can be determined by taking the $1-\alpha$ quantile value of this sampled set.

##### S1.3 Exponentiated gradient updates

The derivative $\nabla\mathcal{L}(\boldsymbol{\pi})$ is as follows:

$$
\begin{aligned}
[\nabla\mathcal{L}(\boldsymbol{\pi})]^{(0)} &= -\sum_{k=1}^{K} g_k(\pi^{(0)})\mathbf{1}_{(s_k^{(0)},s_k^{(1)})=(1,0)} \\
[\nabla\mathcal{L}(\boldsymbol{\pi})]^{(1)} &= -\sum_{k=1}^{K} g_k(\pi^{(1)})\mathbf{1}_{(s_k^{(0)},s_k^{(1)})=(0,1)} = 0
\end{aligned}
\tag{S3}
$$

where

$$
\begin{aligned}
g_k(x) &= \frac{\sum_{i=1}^{n_k}\sum_{j=0}^{\min(m_k,i)} c_{i,j,k}\cdot[ix^{i-1}(1-x)^{n_k-i}-(n_k-i)x^i(1-x)^{n_k-i-1}]}{\sum_{i=0}^{n_k}\sum_{j=0}^{\min(m_k,i)} c_{i,j,k}\cdot x^i(1-x)^{n_k-i}} \\
&= \frac{\sum_{i=1}^{n_k}\sum_{j=0}^{\min(m_k,i)} c_{i,j,k}\cdot(i-xn_k)x^{i-1}(1-x)^{n_k-i-1}}{\sum_{i=0}^{n_k}\sum_{j=0}^{\min(m_k,i)} c_{i,j,k}\cdot x^i(1-x)^{n_k-i}},
\end{aligned}
\tag{S4}
$$

with $c_{i,j,k} = \binom{n_k}{i}\cdot p_m(i,j)p_u(n_k-i, m_k-j)$.

### S2 Supplementary results

#### S2.1 Evaluation of the Smallwood method with a 2-kb window size

To further compare the various VMR-detection methods using similar window sizes, we attempted to run Smallwood and scMET with the same bandwidth/window size as vmrseq and scbs, i.e., 2,000 bp. However, scMET encountered computational errors with smaller window sizes, limiting our evaluation to Smallwood using 2,000-bp windows. The default step size in Smallwood (600 bp) was maintained throughout.

Fig. S24-S27 present comparisons between two window sizes applied to the Smallwood method in synthetic benchmarking experiments. While slight improvements were observed in performance metrics with the 2000-bp window, these gains were marginal, and Smallwood still did not achieve a comparable level of performance as vmrseq or scbs. A similar conclusion can be made for the same comparison applied to the mouse frontal cortex dataset from Luo et al. [1] (Fig. S28). Specifically, while the 2000-bp windows improved the nearest neighbor count score when all selected VMRs were included, overall performance across both window sizes remained similar and fell short of matching the accuracy of vmrseq and scbs.

#### S2.2 Evaluation of clustering performance using the Silhouette score

In addition to the nearest neighbor count score, the region detection methods were also evaluated using an average Silhouette score across cells [6]. Specifically, the scores were computed based on the same cell-to-cell dissimilarity matrices used for computing the nearest neighbor scores in Fig. 3b ("Methods"). The function `silhouette` in R package `cluster` (version 2.1.3) was used for computing the Silhouette scores.

Fig. S29 illustrate the clustering performance evaluated with Silhouette score. It demonstrates a similar message as Fig. 3b, where the regions detected by vmrseq exceed the ones from alternative methods. However, we notice that the Silhouette scores are relatively low despite the UMAP figures (Fig. S6) indicating a decent clustering performance. This is presumably due to the 'curse of dimensionality' [7, 8], where high-dimensional representation results in little distinctions between the maximum and minimum distances between data points. Compared to the nearest neighbor count score which solely depends on the rank of the distances, the Silhouette score is more susceptible to the high dimensions of numerical vectors (i.e., the vectors of regional average methylation) representing each cell since it is computed directly based on the cell-to-cell distances.

#### S2.3 Evaluation of clustering performance with dimension reduction

SINBAD [9] employs a methodology that performs dimensionality reduction on large bins across the genome without any feature selection before cell clustering. We conducted a series of experiments to compare the efficacy of various region selection methods against a baseline approach similar to SINBAD. In this baseline approach, principal component analysis (PCA) was employed for dimensionality reduction on genome-wide 100 kb bins. To ensure a fair and unbiased comparison that mitigates the effects of the curse of dimensionality [7, 8], PCA was consistently applied to the $n_{region} \times n_{cell}$ matrix of regional average methylation levels before proceeding with the comparison.

The imputation scheme in SINBAD was adopted to support PCA computation, which uses the population mean to replace the missing values in the methylation level matrix. The loading matrix of the top 10 principal components (PCs) explaining the most variance was used for computing the cell-to-cell dissimilarity distance. We used the same distance metric as in the "Methods" section to compute the dissimilarity matrices, i.e., Manhattan distances between the regional average methylation vectors of each pair of cells. The function `prcomp` in R package `stats` (version 4.2.0) was used for solving PCA.

Fig. S30-S31 displays the results of nearest neighbor count scores and Silhouette scores respectively. With the entire set of detected regions included, all methods achieved clustering performance comparable to the baseline approach. The Silhouette scores of all methods are improved by a considerable margin owing to the dimension reduction step (comparing S31 to Fig. S29). Also notably, dimension reduction significantly enhanced the separation of cell subtypes for both scMET and Smallwood (comparing S30 to Fig. 3b in the main text). These findings underscore the efficiency of the region selection methods, as they are able to maintain high clustering accuracy with a greatly reduced feature set.

#### S2.4 Sensitivity analysis on hyperparameters

Fig. S32 illustrates the results of a sensitivity analysis performed using the Luo et al. [1] dataset to assess the robustness of the nearest neighbor count score with respect to two hyperparameters: the $\alpha$ value and the minimum number of CpGs in the detected regions. We experimented with $\alpha = 0.01, 0.025, 0.05, 0.1$ as they are common choices of the level of significance in hypothesis testing. The minimum number of CpGs were applied with values less or equal to 20 since we would like to capture fine-scale epigenetic changes that may distinguish cell types in this study. Both variably methylated regions (VMRs) and candidate regions (CRs) were evaluated. Similar to the case study on Luo et al. [1] data in main text, two cell type labeling criteria, the broad classes and the subtypes were used for evaluation.

When the cells are labeled by the broad classes (i.e., excitatory and inhibitory), clustering performance of the methods is particularly robust to hyperparameters (Fig. S32a). On the other hand, the nearest neighbor count score with subtype labels exhibits variation across settings of hyperparameters (Fig. S32b). A higher $\alpha$ value induced a larger number of CpG loci included in detected regions, while the increase of minimum CpG count caused a decrease of detected loci (Fig. S32c-f). Among possible parameter pairs, $\alpha = 0.05$ and minimum CpG count $= 5$ are shown to be a reasonable choice for optimal inference of detecting VMRs.

### S3 Supplementary figures

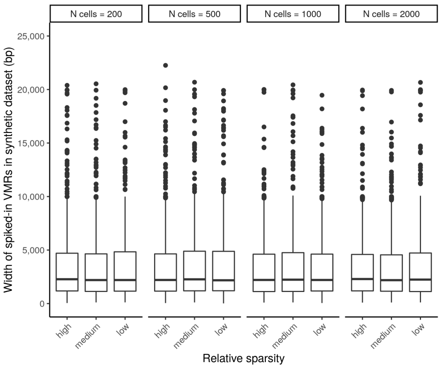


**Fig. S1** Distribution of width of spiked-in VMRs in synthetic datasets used for simulation studies. Width is evaluated in base pairs. Panels are partitioned by the number of subsampled cells and x-axis indicates the sparity level. Note that all settings of 'number of subpopulations' (i.e., $\rho \in \{2, 3, 4, 5, 8, 12, 20\}$) uses identical set of spiked-in VMRs so share the same width distribution.

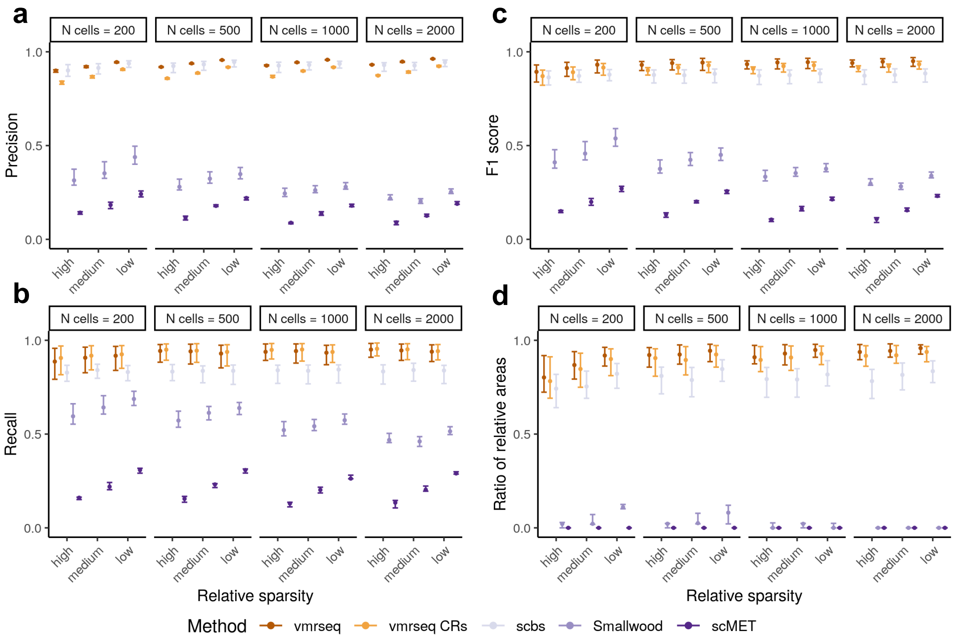


**Fig. S2** *CpG-based* metrics evaluated on *simulated VMRs*, including **a** precision, **b** recall, **c** F1 score and **d** ratio of relative areas ("Methods" of the main text). Each interval consists of points originated from different number of subpopulations. Dot and boundaries of each interval indicates the maximum, median and minimum value of metric. Recall, precision and F1 score are computed using default parameter setting in each method.

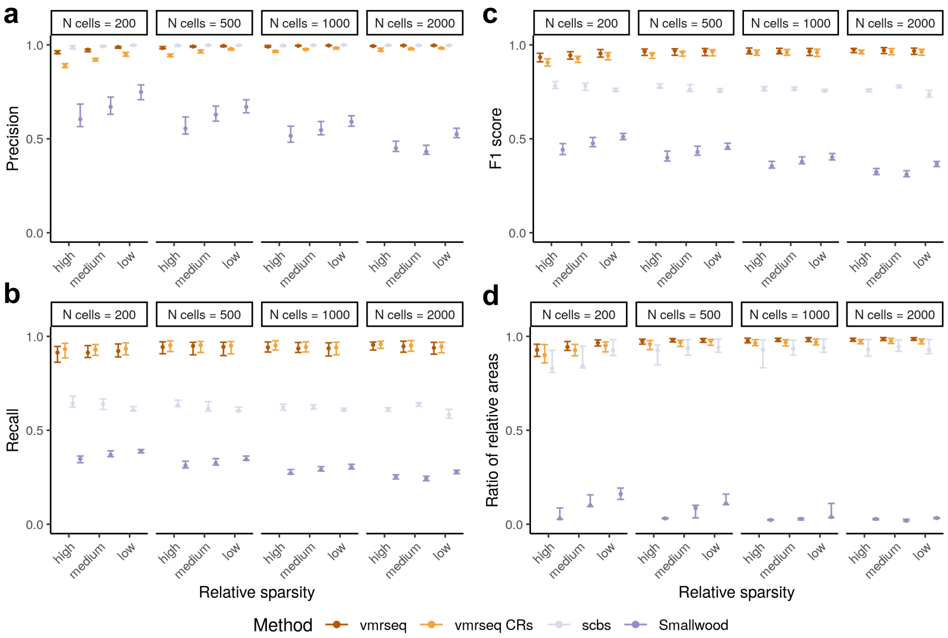


**Fig. S3** *Region-based* metrics evaluated on *synthetic chromosomes*, including **a** precision, **b** recall, **c** F1 score and **d** ratio of relative areas ("Methods" of the main text). A region-level true positive is defined as at least 3 sites overlap between detected and true VMR. Each interval consists of points originated from different number of subpopulations. Dot and boundaries of each interval indicates the maximum, median and minimum value of metric. Recall, precision and F1 score are computed using default parameter setting in each method.

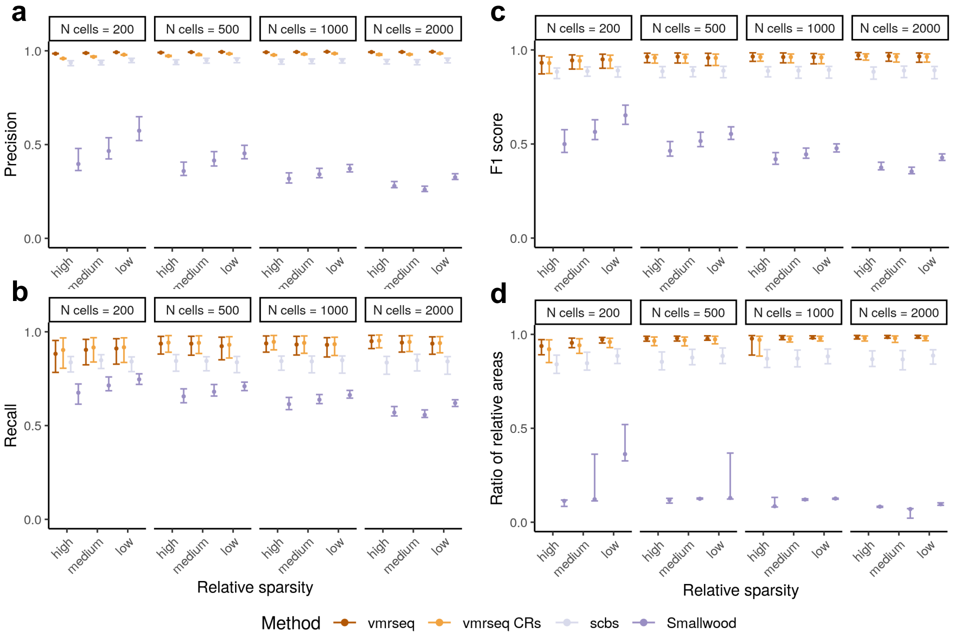


**Fig. S4** *CpG-based* metrics evaluated on *synthetic chromosomes*, including **a** precision, **b** recall, **c** F1 score and **d** ratio of relative areas ("Methods" of the main text). Each interval consists of points originated from different number of subpopulations. Dot and boundaries of each interval indicates the maximum, median and minimum value of metric. Recall, precision and F1 score are computed using default parameter setting in each method.

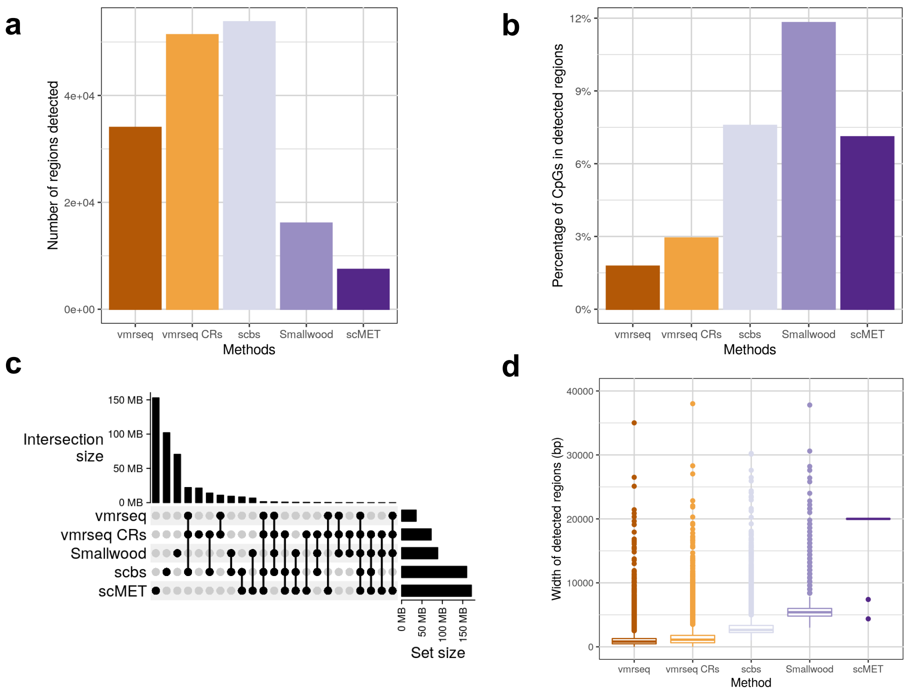


**Fig. S5** Basic output summary of various methods applied on Luo et al. [1] data. For Smallwood, detected windows that overlap are merged and count as one region. **a** Total number of regions detected by each method. **b** Percentage of CpG sites out of total covered sites in regions detected by each method. **c** UpSet plot showing intersection sizes of detected regions from the methods in base pair units. **d** Distribution of region width detected by each methods in the unit of base pair.

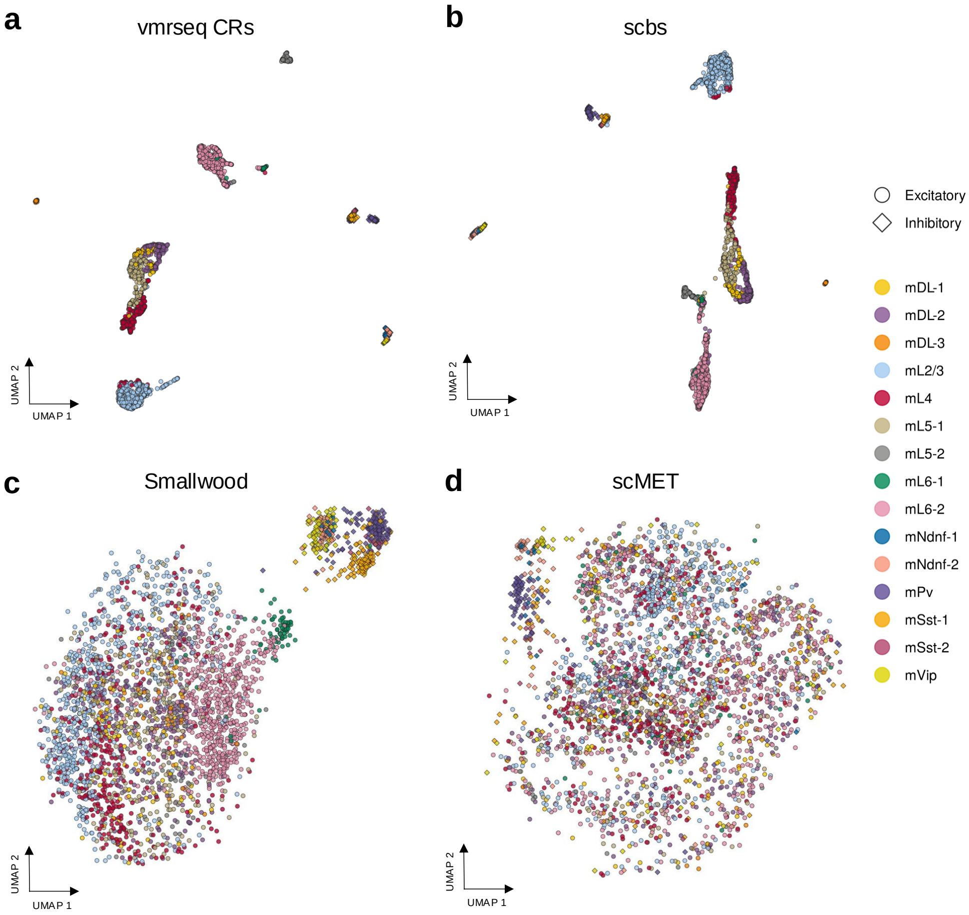


**Fig. S6** UMAP visualization of neuron subpopulations annotated in the Luo et al. [1] dataset for methods other than vmrseq: **a** candidate regions, **b** scbs, **c** Smallwood and **d** scMET. See Fig. 3a for UMAP visulization of vmrseq. For each method, UMAP coordinates were computed from cell-to-cell dissimilarity matrix based on regional average methylation levels of detected regions ("Methods" of the main text). Shape indicates broad cell classes and color indicates the subtypes.

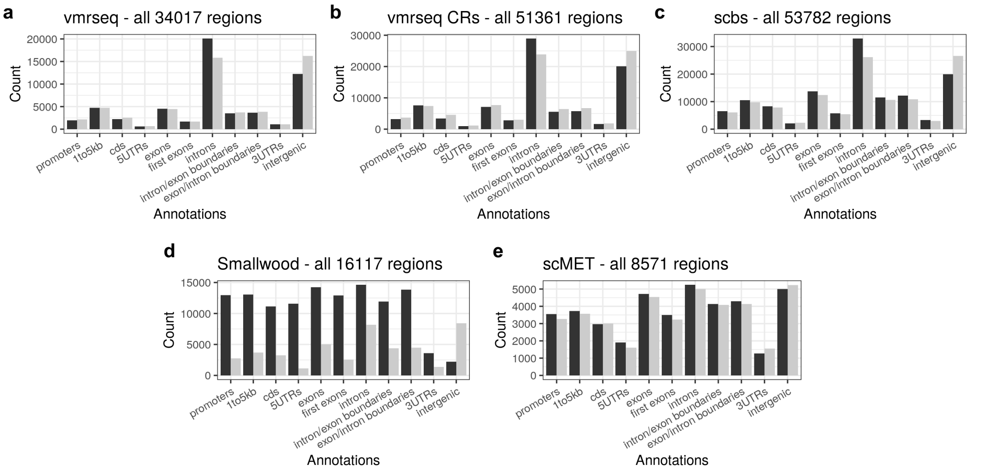


**Fig. S7** Summary of gene context annotation of regions detected in the Luo et al. [1] dataset by **a** vmrseq **b** candidate regions, **c** scbs, **d** Smallwood and **e** scMET. Black bars represent number of detected regions overlapping various types of gene contexts; grey bars represent number of randomized regions overlapping the gene contexts. See "Methods" of the main text for details of the implementation.

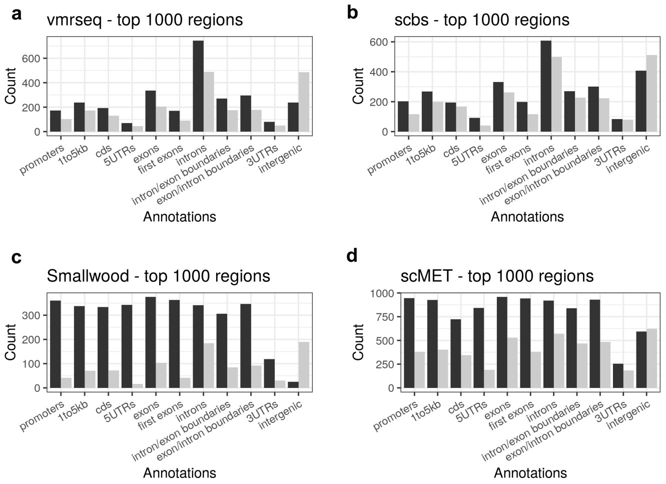


**Fig. S8** Summary of gene context annotation of top 1000 regions from **a** vmrseq **b** scbs, **c** Smallwood and **d** scMET detected in the Luo et al. [1] dataset. Category 'candidate regions' is absent due to the lack of ranking. Black bars represent number of detected regions overlapping various types of gene contexts; grey bars represent number of randomized regions overlapping the gene contexts. See "Methods" of the main text for details of the implementation.

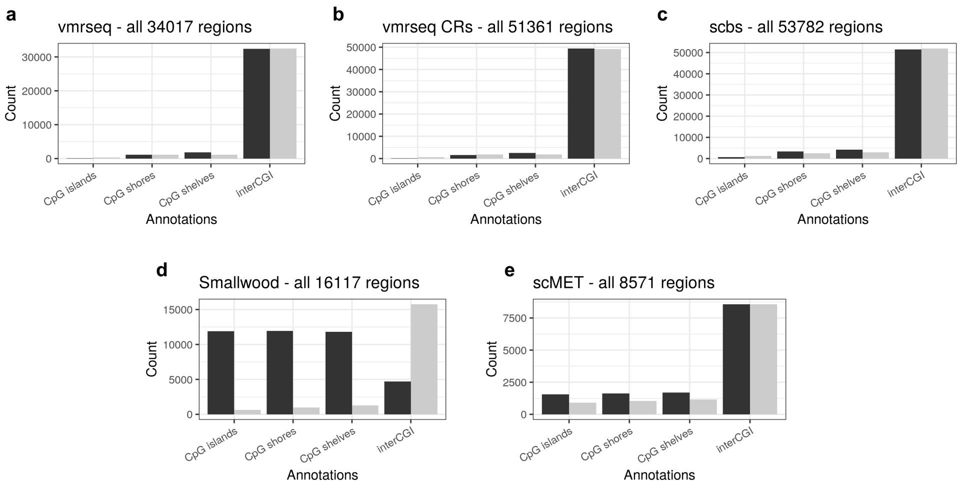


**Fig. S9** Summary of CpG region type annotation of regions detected in the Luo et al. [1] dataset by **a** vmrseq **b** candidate regions, **c** scbs, **d** Smallwood and **e** scMET. Black bars represent number of detected regions overlapping various types of CpG regions; grey bars represent number of randomized regions overlapping the CpG region types. See "Methods" of the main text for details of the implementation.

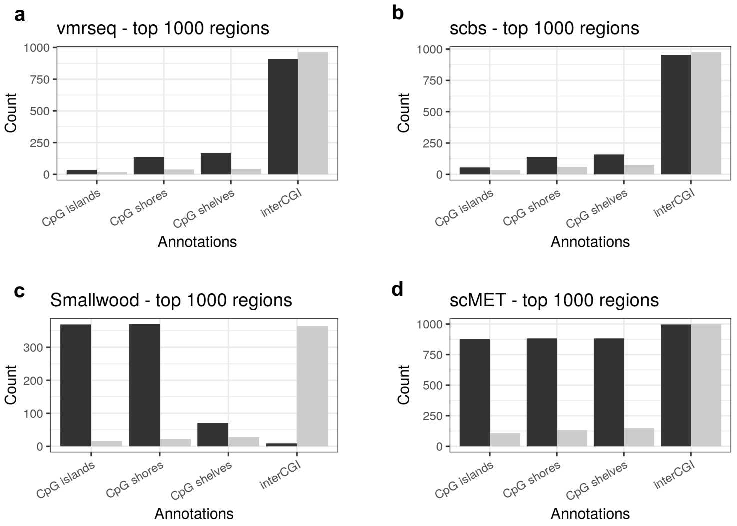


**Fig. S10** Summary of CpG region type annotation of top 1000 regions from **a** vmrseq **b** scbs, **c** Smallwood and **d** scMET detected in the Luo et al. [1] dataset. Category 'candidate regions' is absent due to the lack of ranking. Black bars represent number of detected regions overlapping various types of CpG region; grey bars represent number of randomized regions overlapping the CpG region types. See "Methods" of the main text for details of the implementation.

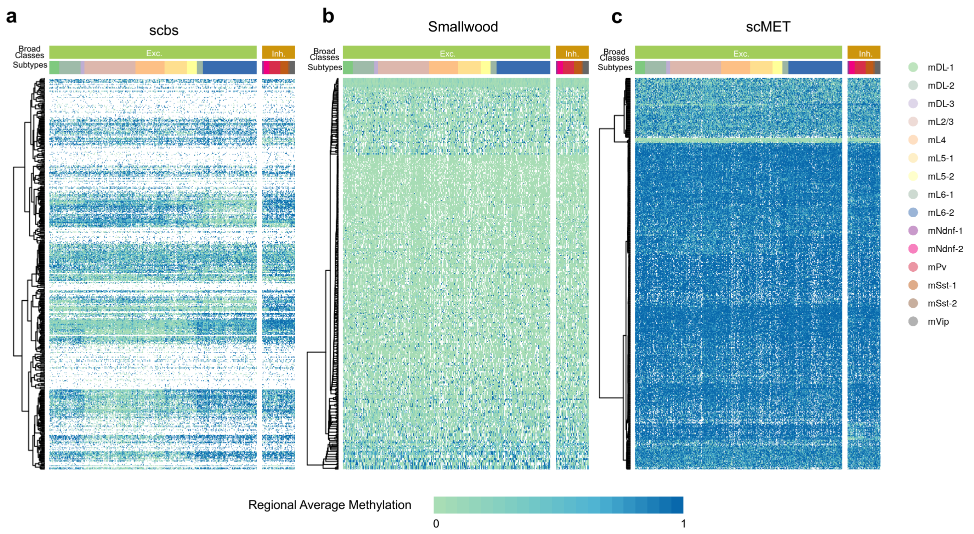


**Fig. S11** Heatmap of regional average methylation level of top-ranked 500 VMRs detected in the Luo et al. [1] dataset by **a** scbs, **b** Smallwood and **c** scMET. Rows are sorted by hierarchical clustering. White color in heatmap represents missing value. Two color bars on top of each heatmap annotate the broad cell classes and subtypes respectively.

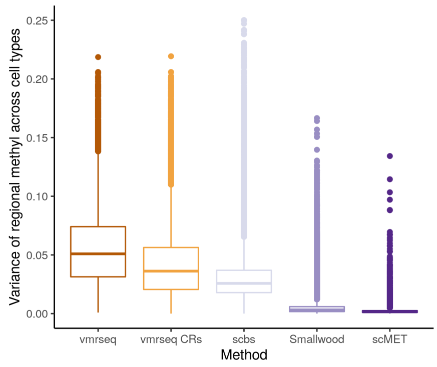


**Fig. S12** Distribution of subtype-to-subtype variance in regional methylation levels computed from detected regions in the Luo et al. [1] dataset by various methods. Regional methylation of a cell subtype is computed by taking the average of regional mean methylation across cells in the subtype. Each data point in a box represents a region.

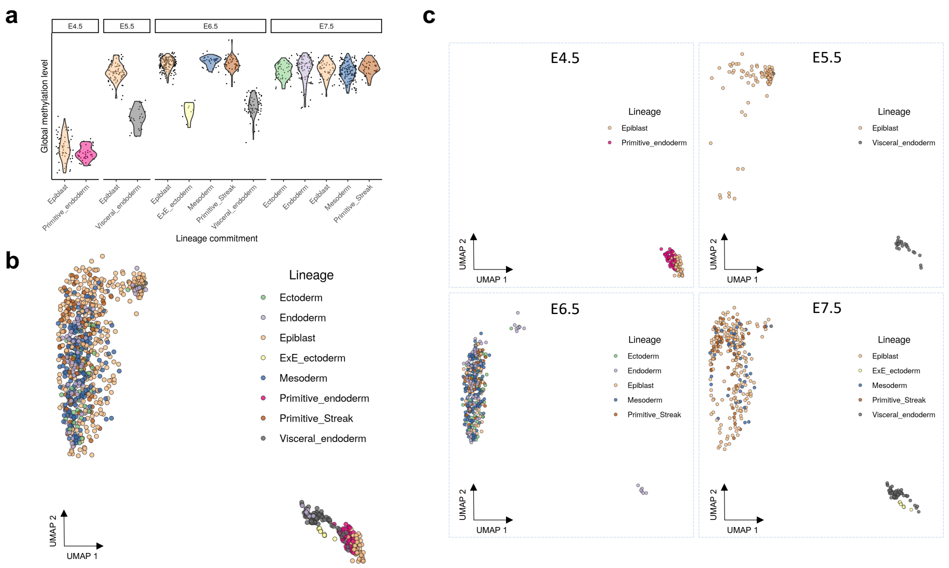


**Fig. S13** Results regarding lineage commitments of the Argelaguet et al. [10] dataset. **a** Distribution of individual-cells global methylation level from the lineage commitments. Each dot represents a cell. Colors represent the lineage commitments. Panels represent developmental stages (E4.5-E7.5) to which the cells belong. **b** UMAP representation with coordinates derived from methylation levels of VMRs output by vmrseq, colored by lineage commitments. **c** UMAP representation in **b** partitioned into panels of developmental stages.

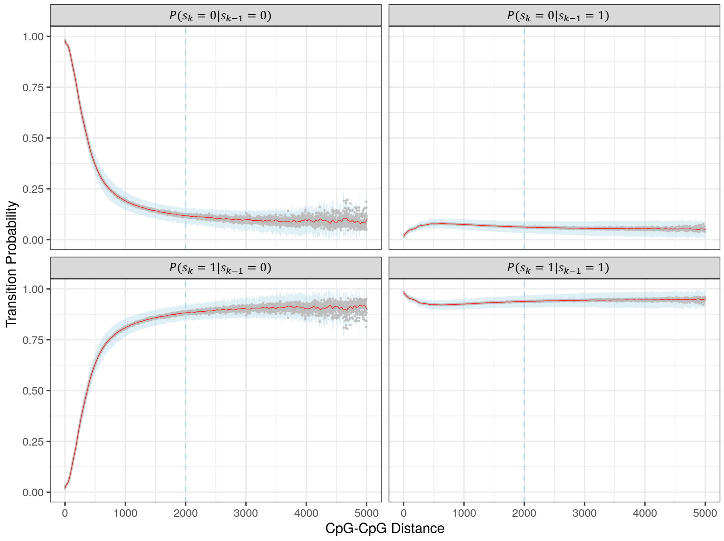


**Fig. S14** Default transition probability distribution with respect to between-CpG distance for one-grouping model. This distribution was trained on aggregated methylation profiles of subtypes. Each panel depicts the probability of a transition mode from one CpG to the next downstream. Grey dots represent estimated transition probability before smoothing; read lines represent the probability after LOESS smoothing. See details for training procedure in section S1.1.2.

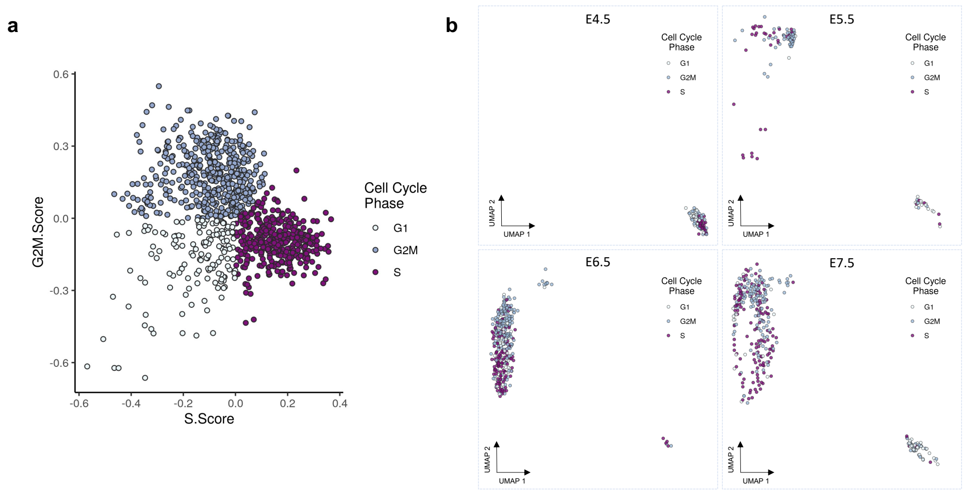


**Fig. S15** Results regarding cell cycle phases of the Argelaguet et al. [10] dataset. **a** Cell cycle phase scores based on canonical markers computed by `CellCycleScoring` function from R pacakge `Seurat` (see "Methods" of the main text for details). Each dot represents a cell, colored by the cell cycle phases annotated by the same function. **b** UMAP representation in Fig. 4b partitioned into panels of developmental stages.

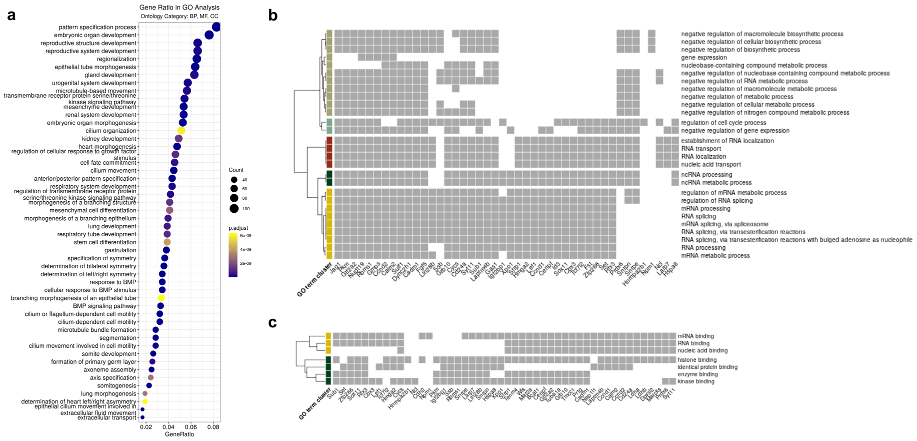


**Fig. S16** Gene ontology (GO) over-representation analyses (ORA) involved in the case study of Argelaguet et al. [10] data. **a** Top 50 significant ontology terms from GO ORA using the list of selected highly variable genes. All three orthogonal ontologies in `clusterProfiler` package, i.e. molecular function (MF), biological process (BP), and cellular component (CC), are included. See "Methods" of the main text for implementation details. **b, c** Gene component of significant ontology terms from subontologies **b** BP and **c** MF, using labeled genes in Fig. 4e. x-axis includes gene symbols found in at least one of the significant terms. Grey squares represent the presence of genes on x-axis in the ontology terms. Hierarchical clusterings of the ontology terms were performed on similarity matrix of the GO terms. The similarity matrices were obtained using `pairwise_termsim` function from R package `enrichplot`.

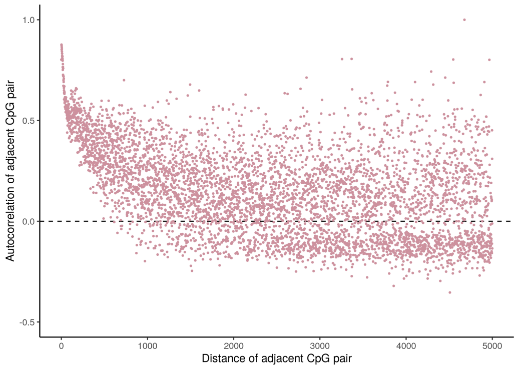


**Fig. S17** Within an individual cell, autocorrelation of DNA methylation between CpGs decreases as genomic distance grows. The cell used here was randomly selected from the Luo et al. [1] dataset. CpG sites with intermediate methylation levels (between 0 and 1) have been removed. Each CpG is paired with its next observed CpG downstream in the same chromosome. X-axis represents the genomic distance (in base pair (bp)) between the CpG pairs. Y-axis represents the Pearson correlation between observed methylation of CpG pairs with identical distance.

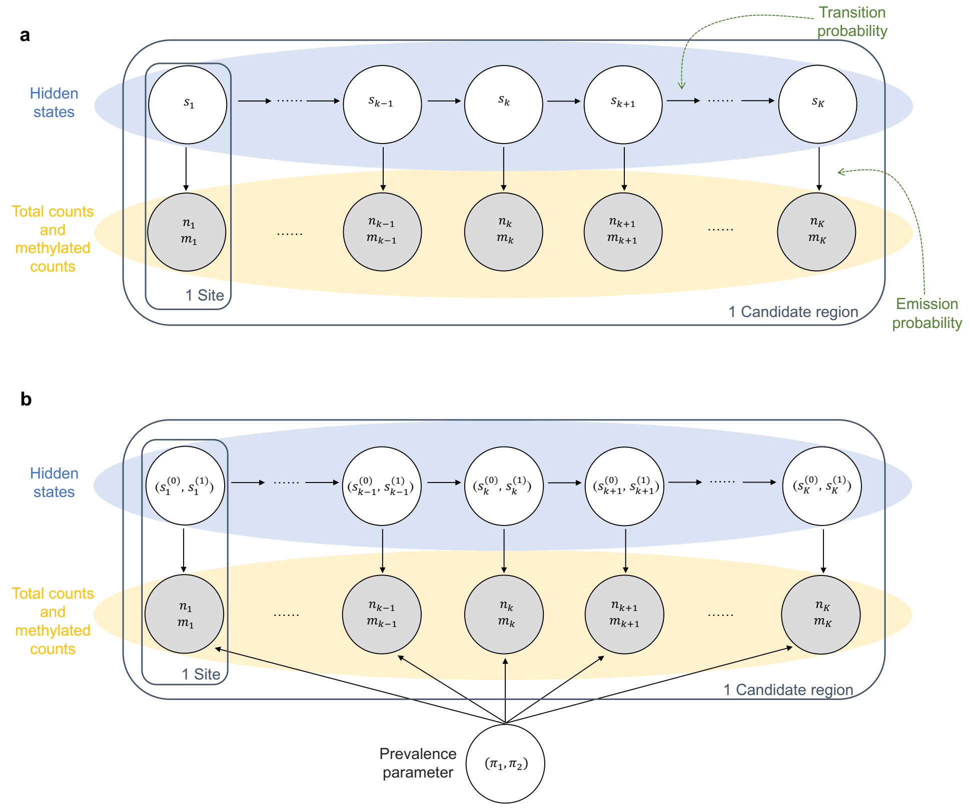


**Fig. S18** Plate notation of **a** one-grouping and **b** two-grouping hidden Markov models respectively. Input values are denoted by gray circles. Model parameters that need optimization are denoted by white circles. Notations are defined in detail in “Methods” of the main text. Note that every candidate region will be modeled independently, thus optimized parameters vary across candidate regions.

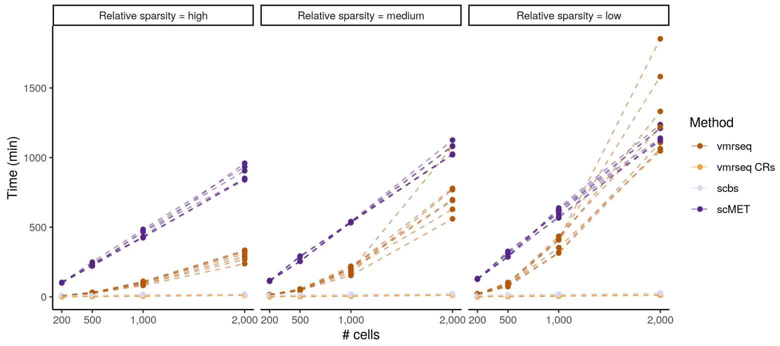


**Fig. S19** Scalability analysis using synthetic data illustrating running times for varying number of cells across methods. Panels represent the sparsity level in synthetic settings. Smallwood was not included in the analysis since the authors did not provide official software. Given a parameter set of time, sparsity and number of cells, seven experiments were run with various numbers of subpopulations. Synthetic data were generated by spiking in simulated VMRs into chromosome 1 of subtype *'IT-L23 Cux1'* from Liu et al. [2] data. Please refer to "Methods" for a detailed description of simulated data generation.

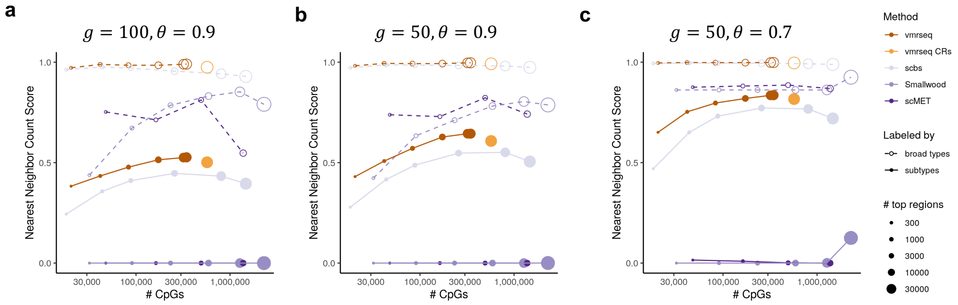


**Fig. S20** **a-c** Nearest neighbor count score with respect to the number of CpG sites in varying numbers of top-ranked VMRs, using alternative choices of parameters $g$ and $\theta$ ("Methods") for clustering performance evaluation of vmrseq and alternative methods on the dataset from Luo et al. [1]. Values of $g$ and $\theta$ are shown on top of each figure. See Fig. 3b for a detailed explanation of the figures.

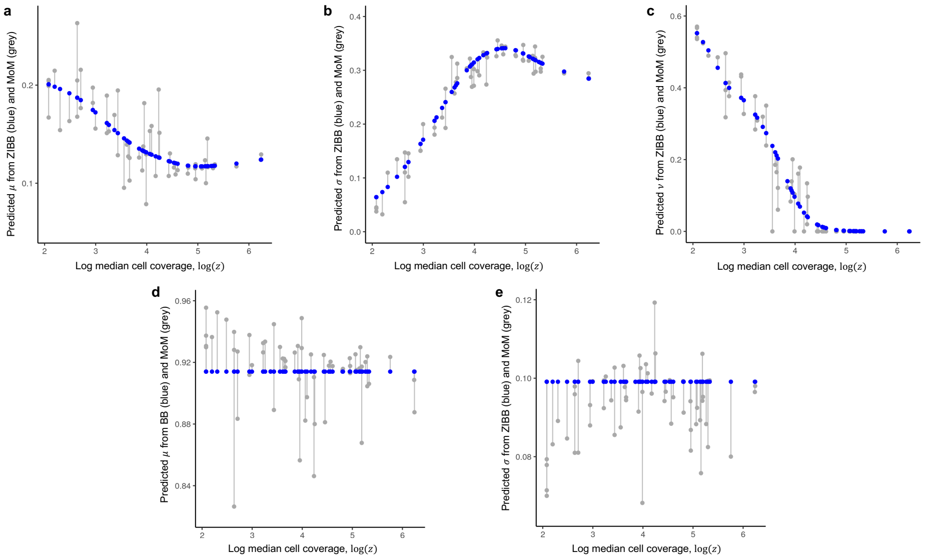


**Fig. S21** Predicted beta prior parameters in emission probability with respect to the coverage level of an input subtype. Blue dots represent predictions from generalized linear regression models trained on all subtypes available in training data; grey dots represent predictions from method of moments (MoM) estimated from individual subtypes separately. **a-c** Prior parameters of $p_U(.,.)$ modeled with zero-inflated beta-binomial (ZIBB) distribution. **b-d** Prior parameters of $p_M(.,.)$ modeled with beta-binomial (BB) distribution. See details for training procedure in section S1.1.3.

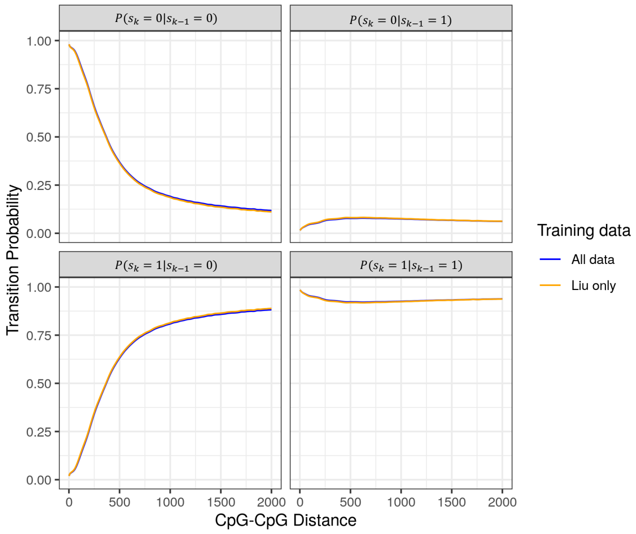


**Fig. S22** Estimates of transition probability distribution before and after the exclusion of Luo et al. [1] data from training data. Same as in Fig. S14, the x-axis represents genomic distance between a pair of CpGs (in base pairs). Blue lines represent original estimates using both Luo et al. [1] and Liu et al. [2] data, and orange lines represent estimates obtained from solely Liu et al. [2] data. See details for training procedure in section S1.1.2.

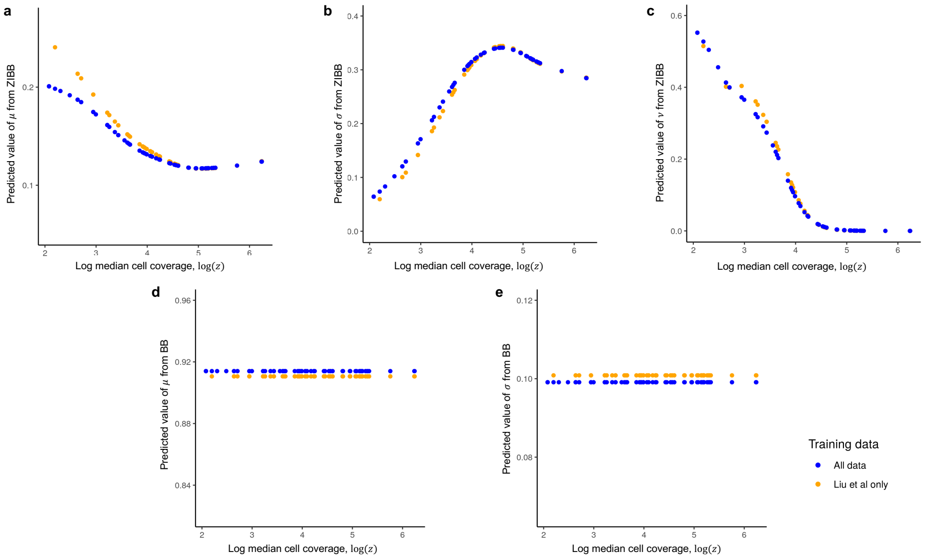


**Fig. S23** Predicted beta prior parameters in emission probability before and after the exclusion of Luo et al. [1] data from training data. Same as in Fig. S21, the x-axis represents the logarithm of the coverage level of an input subtype. Blue dots represent original estimates using both Luo et al. [1] and Liu et al. [2] data, and orange dots represent estimates obtained from solely Liu et al. [2] data. **a-c** Prior parameters of $p_U(.,.)$ modeled with zero-inflated beta-binomial (ZIBB) distribution. **b-d** Prior parameters of $p_M(.,.)$ modeled with beta-binomial (BB) distribution. See details for training procedure in section S1.1.3.

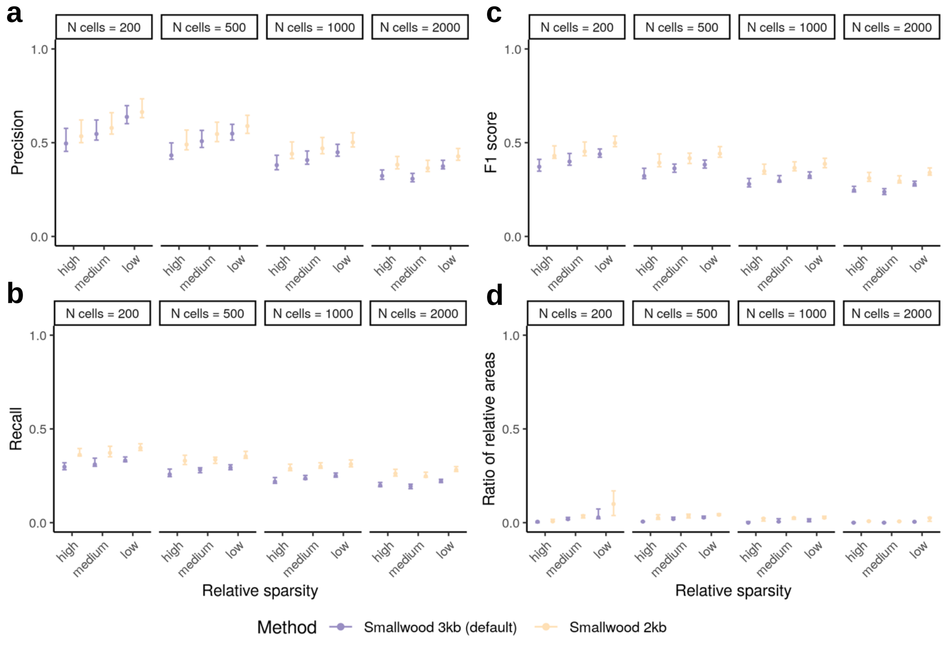


**Fig. S24** Comparison of Smallwood with 3-kb and 2-kb window sizes, evaluated with *region-based* metrics on *simulated VMRs*, including **a** precision, **b** recall, **c** F1 score and **d** ratio of relative areas ("Methods" of the main text). The results in this figure are from the same set of experiments in Fig. 2 of the main text. Same as in the main text, a region-level true positive is defined as at least 3 sites overlap between detected and true VMR. Each interval consists of points originated from different number of subpopulations. Dot and boundaries of each interval indicates the maximum, median and minimum value of metric. Recall, precision and F1 score are computed using default parameter setting in each method.

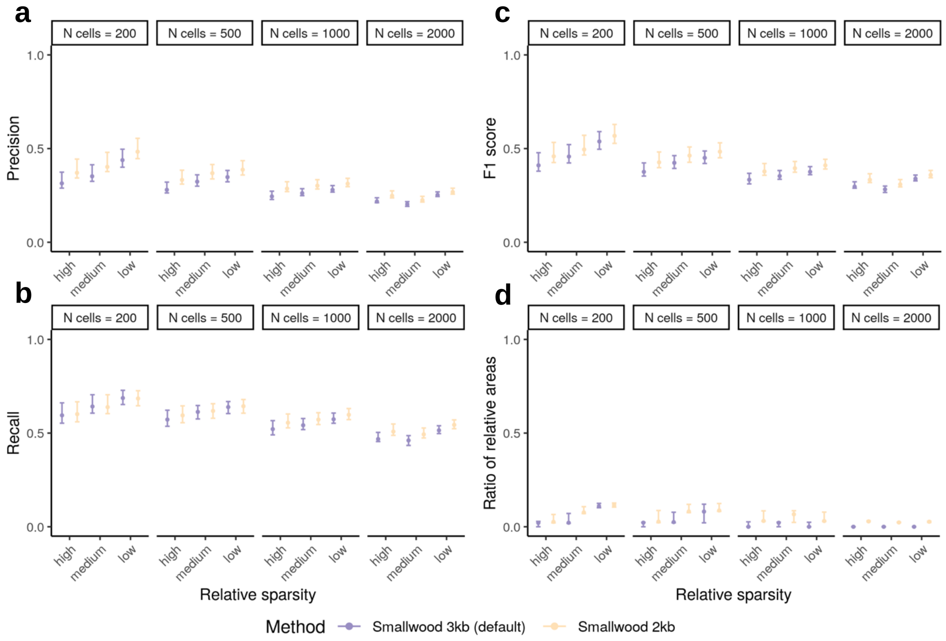


**Fig. S25** Comparison of Smallwood with 3-kb and 2-kb window sizes, evaluated with *CpG-based* metrics on *simulated VMRs*, including **a** precision, **b** recall, **c** F1 score and **d** ratio of relative areas ("Methods" of the main text). The results in this figure are from the same set of experiments in Fig. S2. Each interval consists of points originated from different number of subpopulations. Dot and boundaries of each interval indicates the maximum, median and minimum value of metric. Recall, precision and F1 score are computed using default parameter setting in each method.

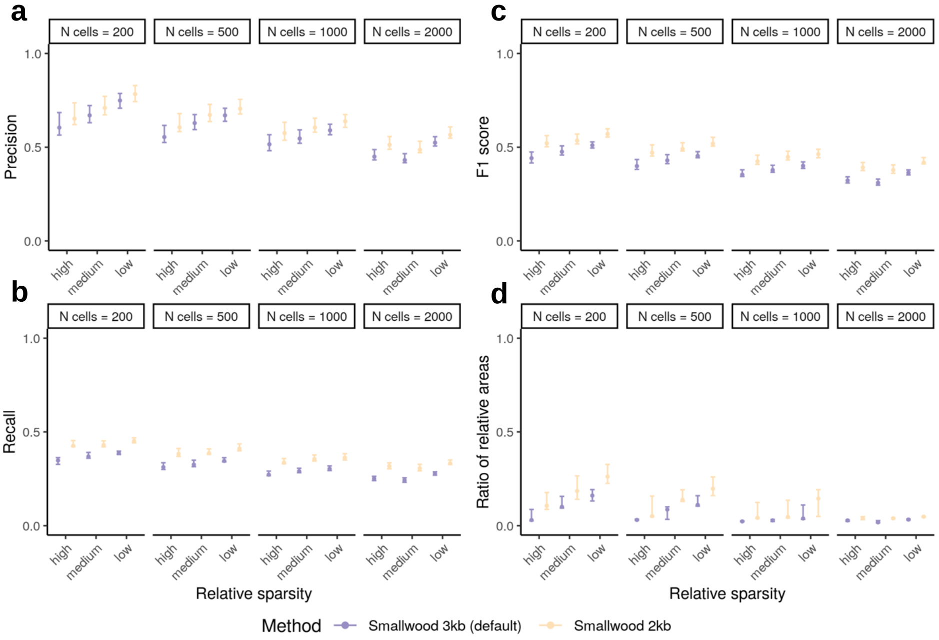


**Fig. S26** Comparison of Smallwood with 3-kb and 2-kb window sizes, evaluated with *region-based* metrics on *synthetic chromosomes*, including **a** precision, **b** recall, **c** F1 score and **d** ratio of relative areas ("Methods" of the main text). The results in this figure are from the same set of experiments in Fig. S3. Same as in the main text, a region-level true positive is defined as at least 3 sites overlap between detected and true VMR. Each interval consists of points originated from different number of subpopulations. Dot and boundaries of each interval indicates the maximum, median and minimum value of metric. Recall, precision and F1 score are computed using default parameter setting in each method.

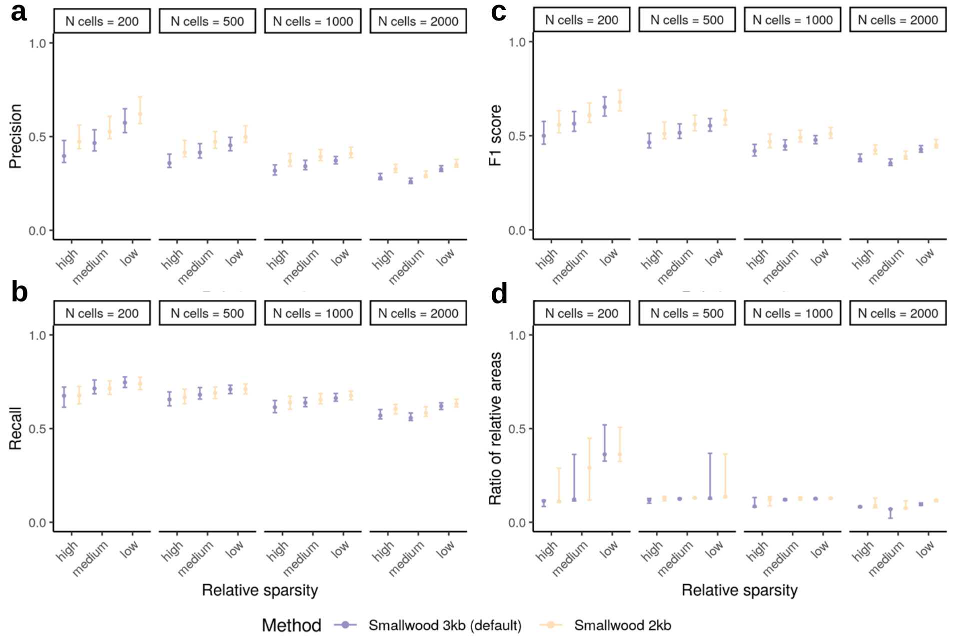


**Fig. S27** Comparison of Smallwood with 3-kb and 2-kb window sizes, evaluated with *CpG-based* metrics on *synthetic chromosomes*, including **a** precision, **b** recall, **c** F1 score and **d** ratio of relative areas ("Methods" of the main text). The results in this figure are from the same set of experiments in Fig. S4. Each interval consists of points originated from different number of subpopulations. Dot and boundaries of each interval indicates the maximum, median and minimum value of metric. Recall, precision and F1 score are computed using default parameter setting in each method.

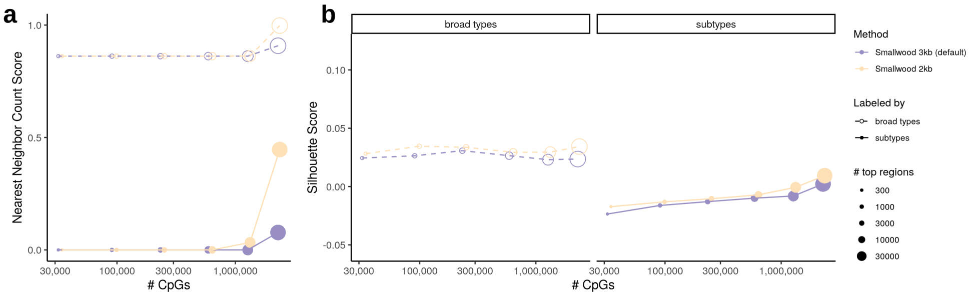


**Fig. S28** Comparison of Smallwood with 3-kb and 2-kb window sizes in the mouse frontal cortex dataset from Luo et al. [1], evaluated with **a** nearest neighbor count score and **b** Silhouette score. The results in this figure are from the same set of experiments in Fig. 3b of main text and Fig. S29. The x-axis is the number of CpG sites in varying numbers of top-ranked VMRs (log-scaled). See Fig. 3b in the main text for a detailed explanation of the legends and how the x-axis is constructed.

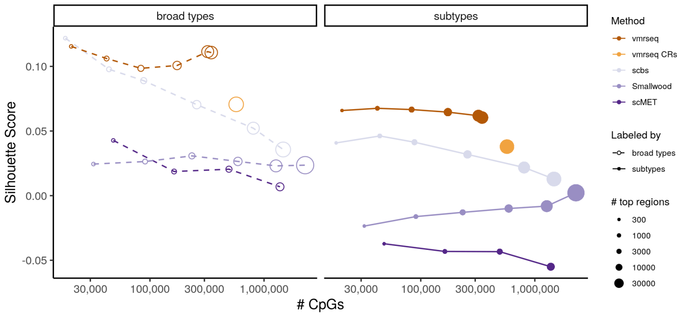


**Fig. S29** Average Silhouette score across cells with respect to the number of CpG sites in varying numbers of top-ranked VMRs, for clustering performance evaluation of vmrseq and alternative methods on the dataset from Luo et al. [1]. See Fig. 3b in the main text for a detailed explanation of the legends and how the x-axis is constructed.

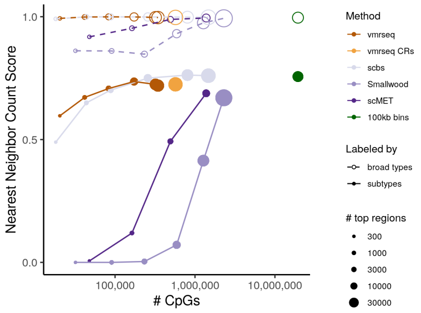


**Fig. S30** Nearest neighbor count score after applying PCA on regional average methylation, with respect to the number of CpG sites in varying numbers of top-ranked VMRs. The scores were based on cell-to-cell dissimilarity matrices computed from PC loadings of the first 10 PCs. Default parameters, $g = 100$ and $\theta = 0.7$, were used for calculation of the nearest neighbor score ("Methods"). See Fig. 3b in the main text for a detailed explanation of the legends and how the x-axis is constructed.

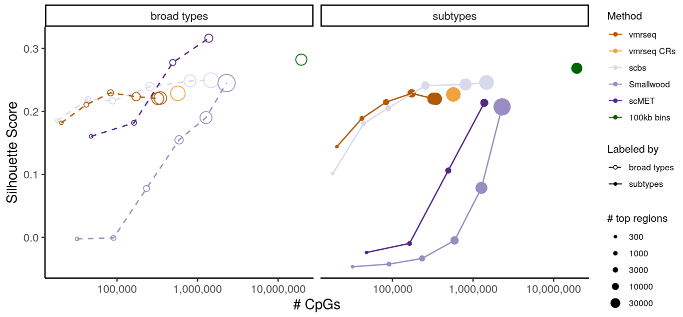


**Fig. S31** Average Silhouette score across cells after applying PCA on regional average methylation, with respect to number of CpG sites in a varying number of top-ranked VMRs. The scores were based on cell-to-cell dissimilarity matrices computed from PC loadings of the first 10 PCs. See Fig. 3b in the main text for a detailed explanation of the legends and how the x-axis is constructed.

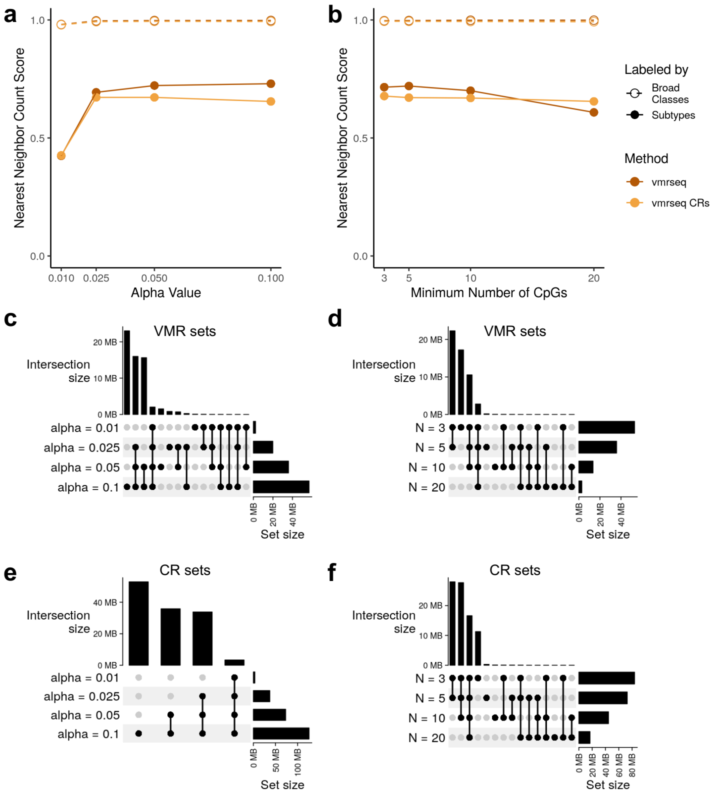


**Fig. S32** Sensitivity analysis on hyperparameters. **a-b** Nearest neighbor count scores evaluated on Luo et al. [1] dataset for various choices of $\alpha$ and minimum CpG count, respectively. Default parameters, $g = 100$ and $\theta = 0.7$, were used for calculation of the nearest neighbor score ("Methods"). For a detailed explanation of the legends, please refer to Fig. 3b in the main text. **c-d** UpSet plot showing intersection of sets of VMR in base pair units from various choices of $\alpha$ and minimum CpG count, respectively. **e-f** UpSet plot showing intersection of set of CR in base pair units from various choices of $\alpha$ and minimum CpG count, respectively.

### Acronyms

*bp* base pair. 23

*CR* candidate region. 8, 35

*PC* principal component. 8
*PCA* principal component analysis. 8

*VMR* variably methylated region. 8, 35

*ZIBB* zero-inflated beta-binomial. 3, 4
